# PI(3)P Signaling by VPS34 Complex II Orchestrates Macropinocytosis

**DOI:** 10.64898/2026.09.03.749235

**Authors:** Astha Neupane, Jared Wollman, Bijaya Pradhan, Jeremiah Donahoe, Joshua Balster, Susmita Poudel, Brandon L. Scott, Adam D. Hoppe, Joel A. Swanson, Natalie W. Thiex

## Abstract

Macrophage macropinocytosis contributes to wound healing, antigen presentation, and resolution of inflammation. Macropinocytosis also facilitates nutrient uptake and growth in macrophages, T cells, and cancer cells. Here, CRISPR/Cas9 whole-genome screens in murine bone-marrow derived macrophages (BMDM) identified genes regulating unstimulated, PMA-, and CSF1-stimulated uptake of the fluorescent pinocytosis solute tracer, Lucifer yellow, to identify novel regulators of macropinocytosis. UVRAG and other members of VPS34 complex II (VPS34-II), which catalyze PI(3)P formation from phosphatidylinositol, were identified as positive regulators of macropinocytosis. Targeted gene disruption of *Uvrag* and *Pik3r3* revealed that VPS34-II is required for efficient macropinocytosis with *Uvrag^sgRNA^* BMDM having fewer but larger macropinosomes. In contrast, depletion of ATG14, a unique component of VPS34 complex I, increased solute uptake and the number of macropinosomes formed per cell. Live-cell imagining of macrophages expressing 2xFYVE-fluorescent protein fusions showed PI(3)P present on the plasma membrane, nascent macropinosomes, and endosomes. The presence of PI(3)P on the plasma membrane prior to macropinocytic cup closure, indicated a novel role for this phosphoinositide species on the plasma membrane. Quantitative imaging of fixed cells shows a decrease in concentration of PI(3)P in *Uvrag^sgRNA^* BMDMs and an increase in concentration of PI(3)P in *Atg14^sgRNA^* BMDMs compared to wildtype BMDMs. Treatment with the VPS34 inhibitor SAR-405 acutely decreased macropinocytosis but maintained AKT phosphorylation suggesting class I PI3K activity and PI(3,4,5)P_3_ production are independent of class III PI3K activity. Overall, these results suggest that PI(3)P is a key phosphoinositide governing macropinocytosis at the plasma membrane and that it is primarily formed via direct phosphorylation of PI rather than via the sequential dephosphorylation of PIP_3_.

## Introduction

Macropinocytosis, or "cell drinking," is a type of actin-dependent fluid-phase uptake where cells non-specifically engulf large amounts of extracellular fluid and solutes. A variety of cells perform macropinocytosis including immune cells, endothelial cells, fibroblasts, and cancer cells, supporting beneficial cellular and physiologic functions as well as pathogenic processes and disease states (Buckley & King 2017; Kerr & Teasdale 2009; Marques et al. 2017). Macropinocytosis is central to several macrophage immune functions including wound healing, antigen presentation, and resolution of inflammation (Bloomfield & Kay 2016).

Despite its importance, progress in defining the molecular machinery of macropinocytosis has been limited. This is due in part to the absence of pathway-specific molecular markers or inhibitors and to the widespread use of fluid-phase tracers that are not exclusively internalized through non-receptor-mediated routes. We recently demonstrated that dextran, one of the most used fluid-phase reporters, binds to mannose receptor C-type 1 (MRC1; CD206) and is internalized by MRC1-expressing cells, including macrophages (Montaner et al. 1999; Wollman et al. 2024). Horseradish peroxidase, another frequently used soluble tracer, is likewise a MRC1 ligand (Feinberg et al. 2021; Montaner et al. 1999). Consequently, uptake of these probes can reflect receptor-mediated endocytosis in addition to pinocytosis, complicating the identification of genes that specifically regulate this process. To minimize this confounding effect, here we used Lucifer yellow, a negatively charged, membrane-impermeant fluorescent dye with no known cellular receptor or transporter (Swanson et al. 1985), as a fluid-phase reporter in genome-wide CRISPR/Cas9 screens. This approach allowed us to identify genes whose disruption selectively altered pinocytic uptake and thereby define molecular machinery required for pinocytosis.

Growth factor-stimulated macropinocytosis requires class I phosphoinositide 3-kinase (PI3K) signaling, which generates phosphatidylinositol (3,4,5)-trisphosphate (PIP_3_) at sites of membrane ruffling as well at the forming macropinosome cup (Araki et al. 1996; Pacitto et al. 2017; Quinn et al. 2021; Yoshida et al. 2009). Interestingly, pharmacological inhibition of PI3Ks via LY294002 does not impair membrane ruffling, extension, or cup formation in murine macrophages, but does eliminate the localized accumulation of the 3-phosphoinositides, PIP_3_ and PI(3,4)P_2,_ and produces macropinocytic cups that fail to fully seal and separate from the plasma membrane (Quinn et al. 2021).

Here, we combine genome-wide CRISPR/Cas9 screens, genetic perturbation of primary macrophages, pharmacological perturbation of class I and III PI3Ks, and live-cell fluorescence microscopy to improve understanding of the molecular mechanisms of macropinocytosis in primary macrophages. Our data identify VPS34-II components, including UVRAG, as key positive regulators of macropinocytosis, and reveal an opposing role for ATG14, a member of VPS34 complex I. We further show that PI(3)P is present at the plasma membrane prior to macropinocytic cup closure and is associated with two distinct PI(3)P-positive vesicle populations: large nascent macropinosomes that undergo directed intracellular transport after scission, and smaller peripheral puncta exhibiting confined dynamics. Together, these findings redefine the events leading up to macropinocytic cup closure and establish a competitive balance between VPS34 complexes in regulating macropinocytic intake.

## Results

### CRISPR/Cas9 whole genome screens identifies VPS34 complex II as a positive regulator of macrophage pinocytosis

To identify genes that regulate pinocytosis, we conducted a series of CRISPR/Cas9 whole genome screens in murine bone marrow-derived macrophages (BMDM). The screens identified gene disruptions conferring a loss-of-function or gain-of-function of Lucifer yellow dye uptake. Briefly, BMDMs were transduced with the Brie sgRNA library containing four guides per gene for most genes in the mouse genome, antibiotic selected to remove untransduced cells, and cultured to allow time for protein depletion. Mutant BMDMs were exposed to Lucifer yellow, and low- and high-fluorescence cells reflecting low and high pinocytosis were sorted by flow cytometry. The sgRNA lentiviral inserts were PCR amplified from genomic DNA and sequenced (Figure 1A). To identify genes regulating pinocytosis, we ran comparative uptake screens under two conditions: 1) CSF1 stimulated and 2) PMA stimulated with three replicates each. CSF1 and PMA signal for pinocytosis via distinct signaling pathways. Therefore, this approach allowed identification of unique genes regulating each uptake route as well as core machinery common to both conditions.

**Figure 1.**
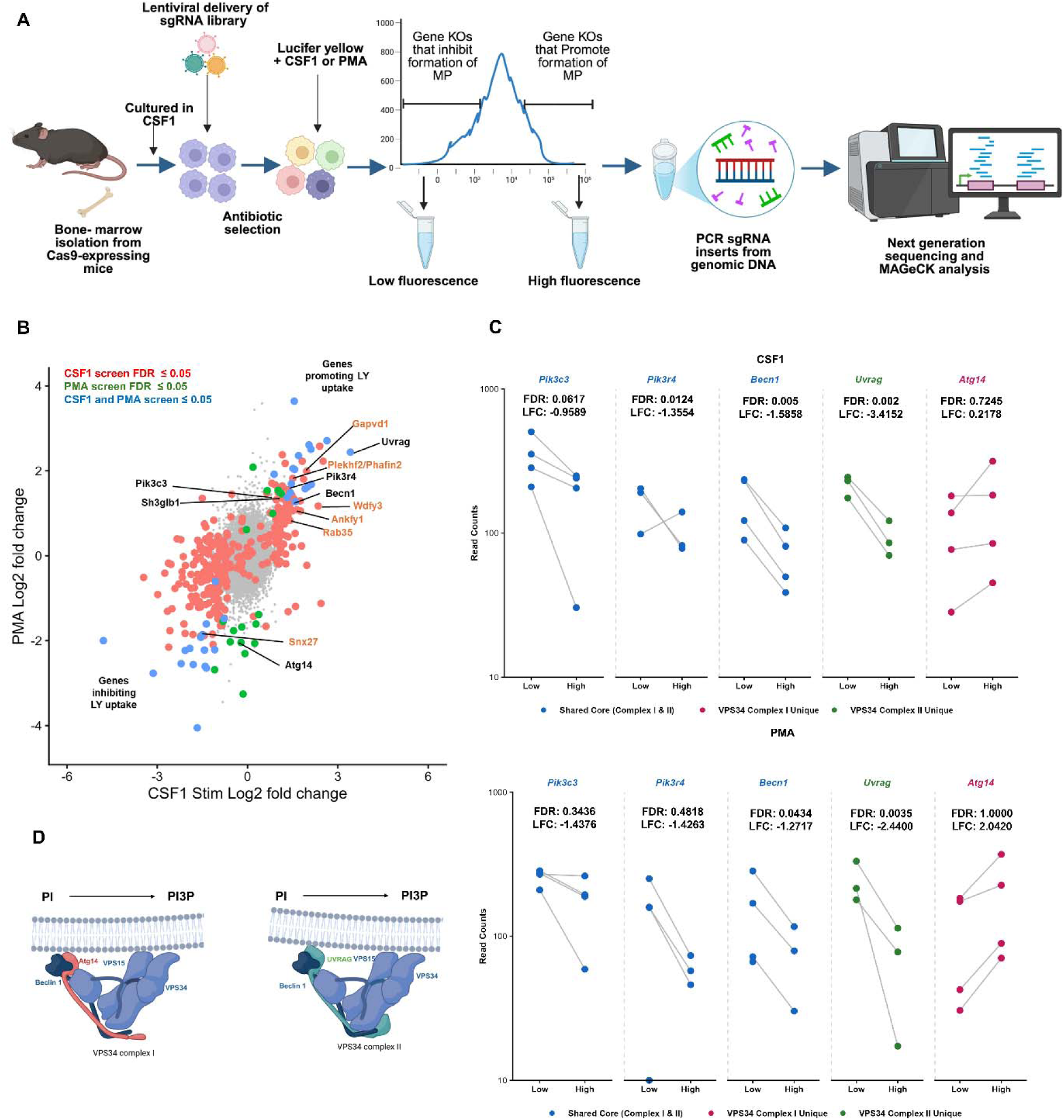
CRISPR/Cas9 whole-genome screens identify VPS34 complex II and effectors of PI(3)P as regulators of macrophage macropinocytosis. A. Model of the CRISPR/Cas9 screen workflow used to identify genes regulating Lucifer yellow uptake in BMDMs. BMDMs isolated from Cas9-expressing mice were transduced with the Brie sgRNA library, selected with antibiotics, and cultured to allow time for gene disruption and protein turnover. Cells were starved of CSF1 overnight, then exposed to 500 mg/ml Lucifer yellow for 30 minutes with CSF1 or PMA. Cells were sorted using flow cytometer, genomic DNA was extracted, sgRNA inserts were amplified and sequenced, and differential enrichment was analyzed using MAGeCK. Image created with BioRender.com. B. Plot highlights the top-ranked screen hits from the CSF1- and PMA-stimulated Lucifer yellow take screens. C. Plots of individual sgRNA read counts from CSF1 and PMA stimulated Lucifer yellow uptake screen replicate 3 for selected genes. D. Schematic structural models of VPS34 complex I and II. Image created with BioRender.com.

The MAGeCK hi-low analysis (Li et al. 2014) was used to calculate RRA score, rank, and log2 fold change (LFC) between the high- and low-fluorescence populations for each individual screen (Supplemental Table 1). The MAGeCK paired analysis (Doench et al. 2016) was performed across biological replicates to generate composite RRA scores, ranks, and LFC. Using a significance threshold of *P* < 0.05, the paired analysis identified positive and negative regulators of Lucifer yellow uptake under each experimental condition. In the CSF1-stimulated screen, 127 genes were identified as positive regulators and 203 as negative regulators of uptake. In the PMA-stimulated screen, 23 genes were identified as positive regulators and 27 as negative regulators (Figure 1B, Supplemental Table 2). Negative LFC values indicate genes whose disruption reduced Lucifer yellow uptake and that therefore function as positive regulators of pinocytic activity, whereas positive LFC values indicate genes whose disruption increased uptake and that therefore function as negative regulators.

Comparative analysis of the CSF1 and PMA screen conditions identified UV radiation resistance associated (*Uvrag*), as a top-ranked hit suggesting it is a component of core pinocytic machinery (Figure 1B-C). Other core members of VPS34-II, *Pik3c3* (VPS34), *Pik3r4* (VPS15), and *Becn1* (Beclin1), were also significantly enriched in the low-drinker population, further validating a role for this complex in macropinocytosis (Figure 1B-C). VPS34-II is a class III PI3-kinase complex that synthesizes PI(3)P from PI and regulates endosomal maturation and endosome-lysosome fusion (Ohashi et al. 2019; Tremel et al. 2021). While <u>class I</u> PI3K have long been studied as regulators of macropinocytosis, the importance of <u>class III</u> PI3Ks has only recently begun to be investigated (Schink et al. 2021). Notably, ATG14, the defining subunit of VPS34 complex I (VPS34-I), exhibited the opposite high-drinker phenotype, suggesting a negative regulatory role in macropinocytosis (Figure 1B-C). Although VPS34-I shares three core subunits (VPS34, VPS15 and Beclin1) with VPS34-II, it incorporates ATG14 instead of UVRAG and is primarily associated with autophagy-related membrane remodeling (Figure 1D) (Ohashi et al. 2019; Tremel et al. 2021).

### UVRAG, but not ATG14, supports PI(3)P production and macropinosome formation

We next used CRISPR/Cas9-mediated targeted gene disruption of individual VPS34 complex I and II components to validate screen hits and to define their specific roles. Gene editing efficiency was confirmed by nanopore sequencing of the sgRNA/Cas9 cut site (Supplemental Figure S1). Consistent with the screen results, loss of core VPS34 components (*Pik3c3*, *Pik3r4*, and *Becn1*) or the VPS34-II-specific subunit UVRAG markedly impaired (∼55%) Lucifer yellow uptake under both CSF1- and PMA-stimulated conditions (Figure 2B-D). Conversely, ATG14 (VPS34-I) depletion caused a significant increase in macropinocytosis (Figure 2B-D). To visualize the phenotypes observed in our flow cytometry, we performed microscopy on BMDMs stimulated with CSF1 and Lucifer yellow. While both *Uvrag^sgRNA^* and *Atg14^sgRNA^* BMDMs maintained robust membrane ruffling (Supplemental Movie 2 and 3, Supplemental figure S2), UVRAG depletion significantly reduced, while ATG14 depletion markedly increased, the number of nascent macropinosomes formed following a CSF1 starve/stim protocol (overnight removal of CSF1 from the medium and subsequent restimulation) (Figure 2D). No differences were observed in the size of formed macropinosomes (Figure 2E).

**Figure 2.**
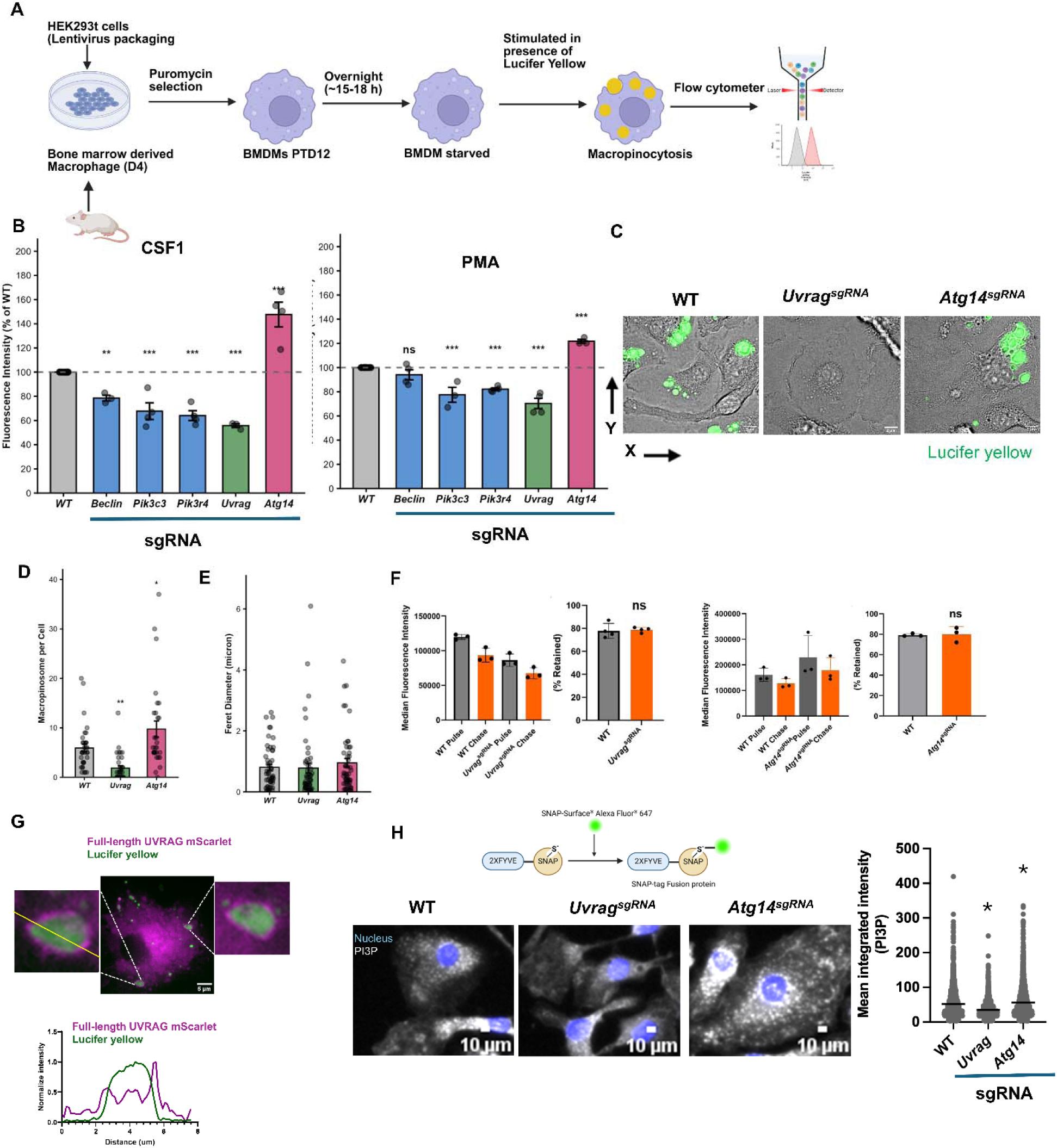
Members of the VPS34 complex II promote efficient macropinocytosis. A. Schematic representation of the isolation of haematopoietic stem cells from mice, their differentiation into macrophages, and lentiviral transduction to generate a specific gene knockout. Image created with BioRender.com. B. Flow cytometry analysis of Lucifer yellow dye (405 nm) in WT and targeted gene knockout in BMDM. BMDMs were starved overnight, stimulated with or without 200 ng/ml CSF1or 100 nM PMA for 30 minutes in the presence of 500 μg/ml Lucifer Yellow, washed and amount of dye uptake was measured by using flow cytometer. Median fluorescence intensity was normalized to WT BMDMs and corrected for autofluorescence using no-dye controls. Statistical analysis was performed using one-way ANOVA followed by Dunnett’s multiple comparisons test. Bar graphs represent the mean ± SD of independent experiments and each dot represents an individual biological replicate. *p < 0.05. Data was analyzed using FlowJo and R. C. BMDMs were starved overnight and stimulated with CSF1 for 5 min in the presence of Lucifer yellow washed and imaged in HBSS using Olympus IX83 microscope, 60X objective. Scale bar 2 um. Representative image for n = 30 cells. D. Quantification of number of feret diameter was done using ImageJ. Dots represent the number of ferret diameter of formed macropinosome. Statistical analysis was performed using GraphPad Prism. Representative image for n = 40 macropinosomes. E. Quantification of number of macropinosome was done using ImageJ. Dots represent the number of macropinosomes. Statistical analysis was performed using GraphPad Prism. Representative image for n = 35 cells. F. BMDMs starved overnight and were exposed to Lucifer yellow in the presence of CSF1 for 90 minutes, and either analyzed immediately in flow cytometer “Pulse” or after a 180-minute chase in DMEM “Chase”. Statistical analysis was performed using one-way ANOVA followed by Tukey multiple comparisons test (95% confidence interval, family-wise alpha = 0.05). Bar graphs represent the mean ± SD of three independent experiments. *p < 0.05. Data was analyzed using FlowJo^TM^ and GraphPad Prism. G. UVRAG regulates micropinocytosis via regulating the synthesis of PI(3)P. Confocal imaging of WT BMDM cells expressing full length Uvrag-mSc stimulated with CSF1 for 2.5 min in the presence of Lucifer yellow, washed and imaged. Representative individual XY optical sections show the localization of Uvrag-mSc (magenta) and Lucifer yellow (green). Scale bar = 5µm. Intensity line-scan analysis of the XY optical section was performed along the yellow line indicated across the vesicle. Relative fluorescence intensities Uvrag-mSc and Lucifer yellow shows localization of Uvrag-mSc around the Lucifer yellow dye. H. High content microscopic imaging BMDMS cultured on coverslips fixed and permeabilized as described in the methods section. Cells were starved overnight and stimulated with CSF1 for 10 minutes. Cells were stained with PI3P-snap-Alexa-647 and Phalloidin. Brightness settings are equal across all images. Scale bar = 10 µm.

Total cellular fluorescence from Lucifer yellow is the amount of dye taken up minus the amount of dye effluxed (Swanson & Watts 1995; Tebeje et al. 2026). Therefore, we sought to determine if these phenotypes were due to changes in macropinocytic uptake or changes in efflux of the Lucifer yellow. BMDM pulsed with Lucifer yellow for 90 min were then chased after washing for 180 min. While less dye was taken up by *Uvrag*^sgRNA^ BMDM and more dye was taken up by *Atg14^sgRNA^* BMDM, the relative amount of dye retained was the same as WT for both (Figure 2F) indicating that VPS34-I and VSP34-II modulate macropinosome formation without contributing significantly to efflux.

### UVRAG regulates macropinocytosis via regulating PI(3)P accumulation on nascent macropinosomes

To further elucidate the mechanism of UVRAG action on macropinosome, we performed live-cell-imaging of full-length UVRAG-mScarlet-I in BMDM cells stably expressing low levels of the fusion protein. Following CSF1 stimulation, Lucifer yellow was readily internalized into macropinosomes, thereby allowing identification of sealed structures. Strikingly, UVRAG was recruited to sealed Lucifer yellow–positive nascent macropinosomes (Figure 2G). As this construct is difficult to express, we did not have enough signal to determine whether it was present on the plasma membrane prior to cup sealing. What we can say is that PI(3)P may be synthesized by VPS34-II directly on nascent macropinosomes.

To delineate the role of UVRAG and ATG14 in regulating production of PI(3)P, we stained UVRAG-deficient, ATG14-deficient and WT BMDM with a recombinant 2xFYVE-SNAP-AF467 probe (Maib et al. 2024). Relative to WT BMDM, *Uvrag^sgRNA^* BMDM contain less PI(3)P and *Atg14^sgRNA^* BMDM contain much higher amounts (Figure 2H). These measurements indicate that UVRAG is essential the majority of cellular PI(3)P synthesis, whereas *Atg14^sgRNA^* is not. These results suggest that PI(3)P production by VPS34-II is crucial for macropinocytosis. VPS34 complex I and complex II compete for shared core subunits (Li et al. 2012), and in the absence of ATG14, we predict that VPS34, VPS15 and Beclin1 are preferentially incorporated into VPS34-II, thus shifting the balance of VPS34 complex assembly toward the UVRAG-containing complex and presumably changing the cellular locations where PI(3)P is synthesized. This is consistent with previous observations in HEK293T cells, where shRNA-mediated ATG14 depletion had limited effects on endomembrane trafficking (Pavlinov et al. 2020). In this model, loss of ATG14 likely favors assembly or activity of UVRAG-containing VPS34-II complex, redirecting PI(3)P to membrane compartments that support macropinocytosis.

### PI(3)P accumulates within macropinocytic cups before sealing and scission

Given that UVRAG was required for both macropinocytosis and PI(3)P production, we hypothesized that VPS34-II may support PI(3)P production or delivery on macropinocytic cups before they seal and detach from the plasma membrane. Macropinocytic cups are open, approximately circular membrane structures generated by the circularization of actin-rich ruffles; subsequent membrane fusion and scission seal the cup and release a nascent macropinosome into the cytoplasm. Macropinocytic cups are difficult to distinguish from macropinosomes without a full three-dimensional volume (Quinn et al. 2021). Therefore, to differentiate open macropinocytic cups from sealed macropinosomes, BMDMs expressing the PI(3)P sensor 2xFYVE-mScarlet-I (2xFYVE-mSc) were stimulated with CSF1, cooled to 4°C and exposed to FM4-64 thereby labeling only plasma membrane that is in contact with the extracellular medium (Maxson et al. 2021; Yoshida et al. 2009). Using this approach, open macropinocytic cups were identified as circular FM4-64-positive structures. Visualization of 2xFYVE-mScarlet-I recruitment revealed that PI(3)P accumulates at the cup stage prior to sealing and scission (Figure 3B).

**Figure 3.**
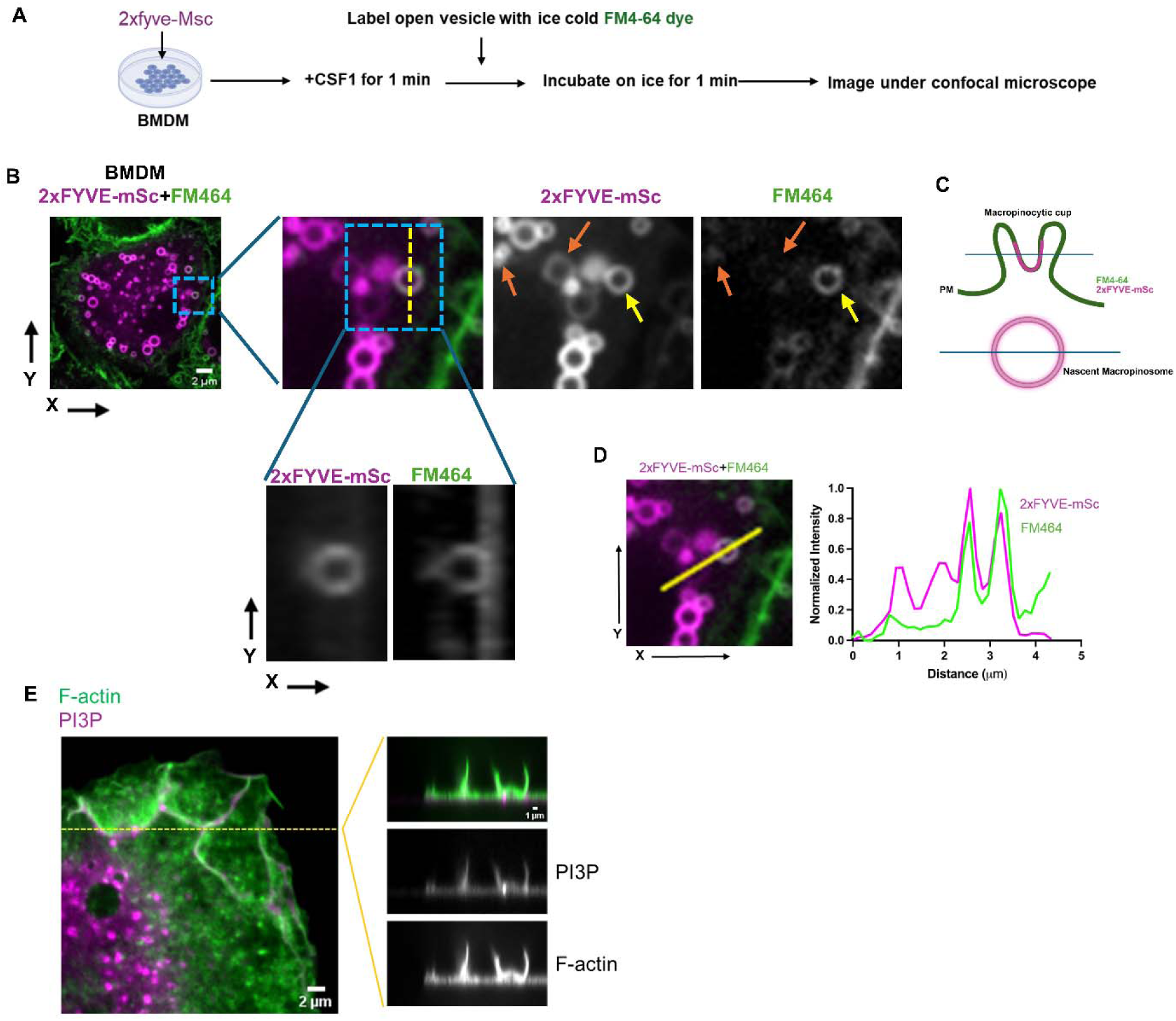
PI(3)P is recruited to the macropinocytic cup following CSF1 stimulation prior to cup closure. A. Confocal imaging of WT BMDMs expressing 2xFYVE-mSc (magenta) stimulated with CSF1 for 1 min. Cells were rapidly cooled and visualized by confocal microscopy. Surface membranes and open macropinosomes were labeled with FM4-64 (green) (see Materials and Methods). Representative individual XY optical sections show the localization of 2xFYVE and FM4-64 in open macropinosomes (yellow arrow) and closed macropinosomes (orange arrow). B. The dashed line across the XY section indicates the position of the corresponding YZ reconstruction. Scale bar = 5 µm. C. Schematic overview of the observed recruitment dynamics from figure A, and B. D. Intensity line-scan analysis of the XY optical section was performed along the yellow line indicated across the vesicle. Relative fluorescence intensities of FM4-64 and 2xFYVE-mScarlet demonstrate their co-localization and recruitment at open macropinosomes. Scale bar = 5 µm. E. Confocal imaging of WT BMDMS cultured on coverslips fixed and permeabilized as described in the methods section. Cells were starved overnight and stimulated with CSF1 for 5 minutes. Cells were stained with PI3P-snap-Alexa-647 and Phallodin. Brightness settings are equal across all images. Scale bar = 10 µm.

Line-scan analysis confirmed overlap of 2xFYVE and FM4-64 signal at the ruffle rim (Figure 3D). We further examined the spatial relationship between PI(3)P and actin-rich membrane ruffles in fixed cells using recombinant 2xFYVE-SNAP-AF647 to detect PI(3)P and phalloidin to label F-actin. PI(3)P-positive puncta were enriched at the base of F-actin-rich ruffles resembling those that give rise to macropinocytic cups (Figure 3E). Together, these observations place PI(3)P within the actin-remodeled plasma membrane domain during cup formation and support a role for PI(3)P, possibly delivered on endosomes to the base of ruffles and cups, before macropinosome closure.

### Class I PI3Ks and VPS34 independently promote macropinocytosis

The appearance of PI(3)P within open cups raised the question of how this lipid is generated. Because sequential dephosphorylation of PIP_3_ can produce PI(3,4)P_2_ and subsequently PI(3)P, one possibility was that the early PI(3)P pool arose downstream of class I PI3K activity. Alternatively, VPS34 could generate PI(3)P directly from phosphatidylinositol. Our previous work showed that LY294002 permits the formation of circular macropinocytic cups but prevents their closure (Quinn et al. 2021). Because LY294002 inhibits both class I PI3Ks and VPS34, that experiment could not distinguish which kinase activity was required. Therefore, we treated BMDMs with 1) a inhibitor cocktail specific to class I PI3Ks, 2) the VPS34-selective inhibitors SAR405 or VPS34-IN1, or 3) LY294002 and measured Lucifer yellow uptake. SAR405 and VPS34-IN1 both greatly reduced Lucifer yellow uptake (Figure 4A) without disrupting AKT phosphorylation (Fig 4B, Supplemental Figure S2 A), indicating that class I PI3Ks and VPS34 promote macropinocytosis through separable signaling pathways.

**Figure 4.**
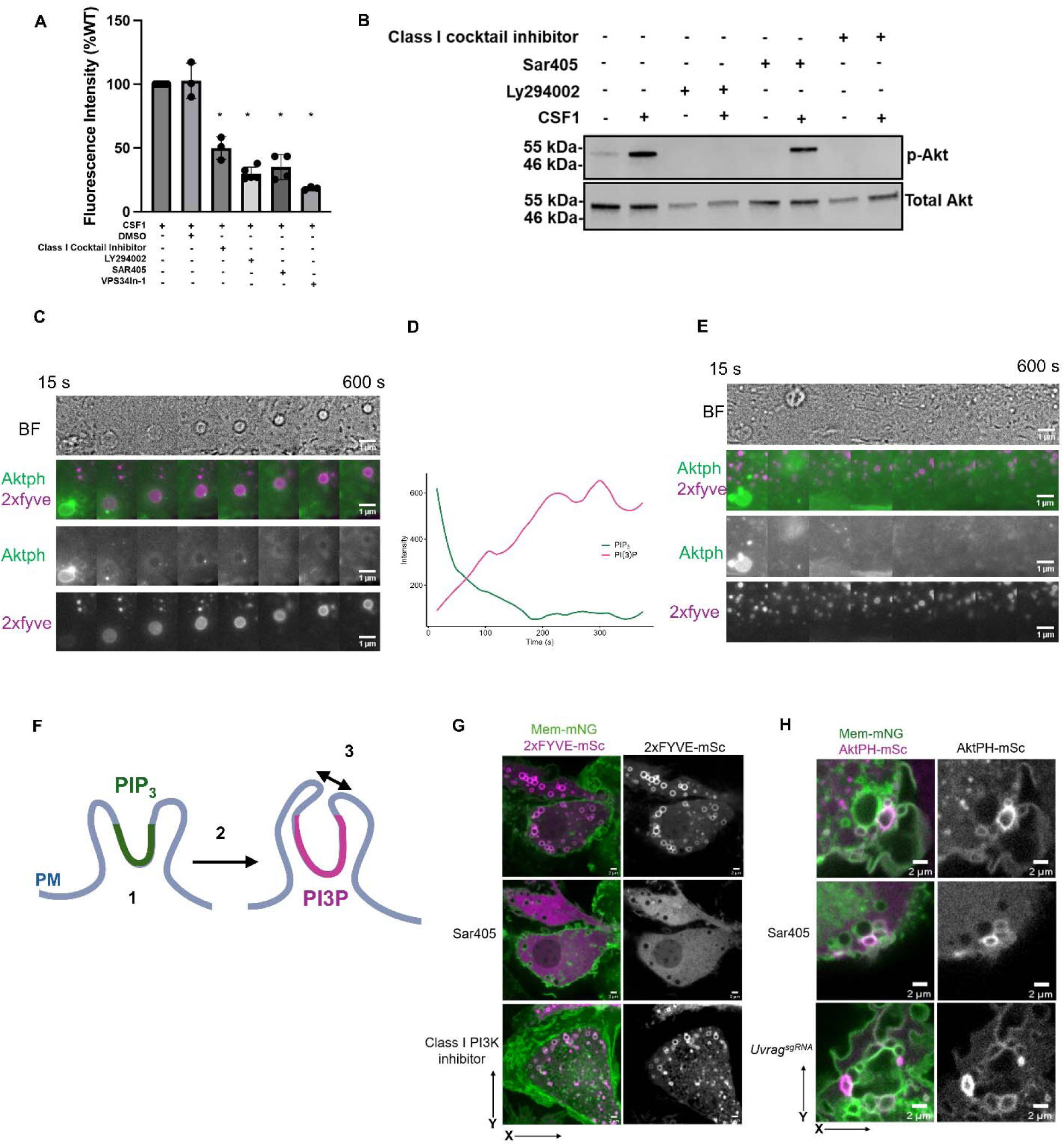
Class I and class III PI3K lipid kinase activity are both required for macropinocytosis, but function independently of each other. (Macropinocytic cups form in the absence of both. A) Flow cytometry analysis of Lucifer yellow dye in WT and drug-treated BMDM (Class I cocktail inhibitor (A66, TGX221, AS252424 and IC87114), Class III kinase inhibitor (SAR-405 and VPS34IN-1) and broad-spectrum PI3K kinase inhibitor (LY294002). BMDMs were starved overnight, treated with drugs for 30 min and stimulated with or without 200 ng/ml CSF1 for 30 minutes in the presence of 500 μg/ml Lucifer Yellow, washed and amount of dye uptake was measured by using flow cytometer. Median fluorescence intensity was normalized to untreated wild-type BMDMs and corrected for autofluorescence using no-dye controls. Statistical analysis was performed using one-way ANOVA followed by Dunnett’s multiple comparisons test (95% confidence interval, family-wise alpha = 0.05). Bar graphs represent the mean ± SD of three independent experiments. *p < 0.05. Data was analyzed using FlowJo and GraphPad Prism. B) Western blot analysis of phosphorylated AKT (p-AKT) and total AKT in WT BMDMs under pharmacological inhibition of PI3K signaling. Cells were treated with a Class I PI3K inhibitor cocktail (A66, TGX221, AS252424, and IC87114), Class III PI3K inhibitors (SAR-405 and VPS34-IN-1), or the broad-spectrum PI3K inhibitor LY294002. BMDMs were serum-starved overnight, pretreated with inhibitors for 30 min, and then stimulated with or without 200 ng/mL CSF1 for 5 min. Cells were subsequently lysed, and protein extracts were resolved by SDS-PAGE and immunoblotted for p-AKT and total AKT. Lysates were derived from the same experimental set but were run on separate gels, which were processed in parallel to ensure comparability across conditions. The data shown are representative of three independent biological replicates performed with similar results. Quantification of p-AKT levels was normalized to total AKT and expressed relative to the CSF1-stimulated control condition (Supplemental S3 A) C) Sequential images of a forming macropinosome in CSF1 stimulated WT BMDMs expressing AktPh-mSc (PIP_3_) and 2xFYVE-mCer (PI3P). WT BMDMs expressing the biosensors were starved overnight and imaged following CSF-1 stimulation. Timestamps represent elapsed time in seconds (15 s frame interval). Representative image for n = 10 macropinosomes/ n=10 cells. Scale bar: 1 µm. See Supplemental Movie 4 and 5. D) Line plot shows corrected fluorescent intensity of PIP_3_( green) and PI3P (magenta) over time. E) Sequential images of failed/unclosed macropinosome in CSF1 stimulated WT BMDMs expressing AktPh-mSc (PIP_3_) and 2xFYVE-mCer (PI3P). WT BMDMs expressing the biosensors were starved overnight and imaged following CSF-1 stimulation. Timestamps represent elapsed time in seconds (15 s frame interval). Representative image for n = 25 macropinosomes. Scale bar: 1 µm. See Supplemental Movie 6 and 7. F) Schematic overview of the observed recruitment dynamics from C, D and E. (1) Open macropinocytic cups initially form at the plasma membrane with local PIP_3_ enrichment (green). (2) PI(3)P (magenta) subsequently accumulates on the cup membrane prior to closure. (3) Recruitment of PI(3)P is required for membrane extension, sealing, and scission (arrow) to form an internalized macropinosome. G) Confocal imaging of BMDMs expressing 2xFYVE-mScarlet (magenta) and Mem-mNG(green), treated with or without PI3K pathway inhibitors and, stimulated with 200 ng/mL CSF1 for 2.5 min. Representative individual XY optical sections show the localization of Akt-PH-mScarlet (magenta) and Mem-mNG(green). The dashed line across the XY section indicates the position of the corresponding XZ or YZ reconstruction. Scale bar = 2 µm. H) Confocal imaging of BMDMs expressing 2xFYVE-mScarlet (magenta) and Mem-mNG(green), treated with or without PI3K pathway inhibitors and, stimulated with 200 ng/mL CSF1 for 2.5 min. Representative individual XY optical sections show the localization of 2xFYVE-mScarlet (magenta) and Mem-mNG(green). The dashed line across the XY section indicates the position of the corresponding XZ or YZ reconstruction. Scale bar = 2 µm.

### PI(3)P accumulation in forming cups does not require class I PI3K activity

To determine whether PI(3)P arrival on macropinosomes is dependent on PIP_3_, we first captured timelapse movies of BMDMs expressing 2xFYVE-mCer and AktPH-mSc (Figure 4C-D, Supplemental Movie 4-5). As previously reported, AktPH-mSc accumulated within membrane ruffles and macropinocytic cups first. As AktPH-mSc signals were diminishing, PI(3)P subsequently appeared as discrete puncta or membrane-associated domains while PIP_3_ remained detectable, demonstrating that the two phosphoinositides can coexist within forming macropinocytic cups and that existing PI(3)P-positive endosomes appeared to partially fuse with the forming macropinosome. We found that PI(3)P recruitment is essential for macropinosome closure and cups lacking PI(3)P failed to mature and regressed back into the plasma membrane (Figure 4E, Supplemental movie 6 and 7, Figure 3F). Among 36 total PIP_3_ positive cups we observed, only the 14 events that acquired PI(3)P successfully matured, while 22 cups lacking PI(3)P aborted and disappeared. To delineate if class I PI3K was required for PI(3)P appearance on open cups, we analyzed FM4-64 labeled open macropinosomes in BMDM treated with the class-I PI3K inhibitor cocktail. Here, we observed efficient recruitment of 2xFYVE-mSc within the open cup (Supplemental Figure S3 G) indicating that VPS34-II can produce PI(3)P on forming macropinosomes independent of class-I PI3K. In addition, treatment of BMDM with the class I PI3K cocktail had minimal effect on the distribution of 2xFYVE-Sc, whereas treatment with Sar405 abolished all endosomal localization (Figure 4G). In comparison, treatment of BMDM with Sar405 or gene disruption of UVRAG did not preclude the localization of AktPH-mSc to forming macropinosomes (Figure 4H, Supplemental Figure S3 C and D, Supplemental movie 8,9 and 10) reinforcing the finding that these two pathways are parallel, and both class I and class III PI3K activities are required for macropinosome formation.

### LLS imaging of shows two distinct classes of PI(3)P positive vesicles

To define how VPS34-II regulates macropinocytosis, we performed lattice light-sheet (LLS) imaging of BMDMs expressing the PI(3)P biosensor 2xFYVE–mScarlet. This high-resolution approach revealed two spatially and morphologically distinct PI(3)P-positive populations: large vesicles corresponding to nascent macropinosomes, and smaller PI(3)P-positive puncta exhibiting centripetal or peripherally restricted movement (Figure 5A; Movie S11). To confirm the identity of the larger vesicle population, cells were stimulated with CSF1 in the presence of Lucifer yellow for 2.5 min prior to imaging. 2xFYVE signal surrounded Lucifer yellow-positive vesicles (Figure 5B–C), confirming that these structures are newly formed macropinosomes.

**Figure 5.**
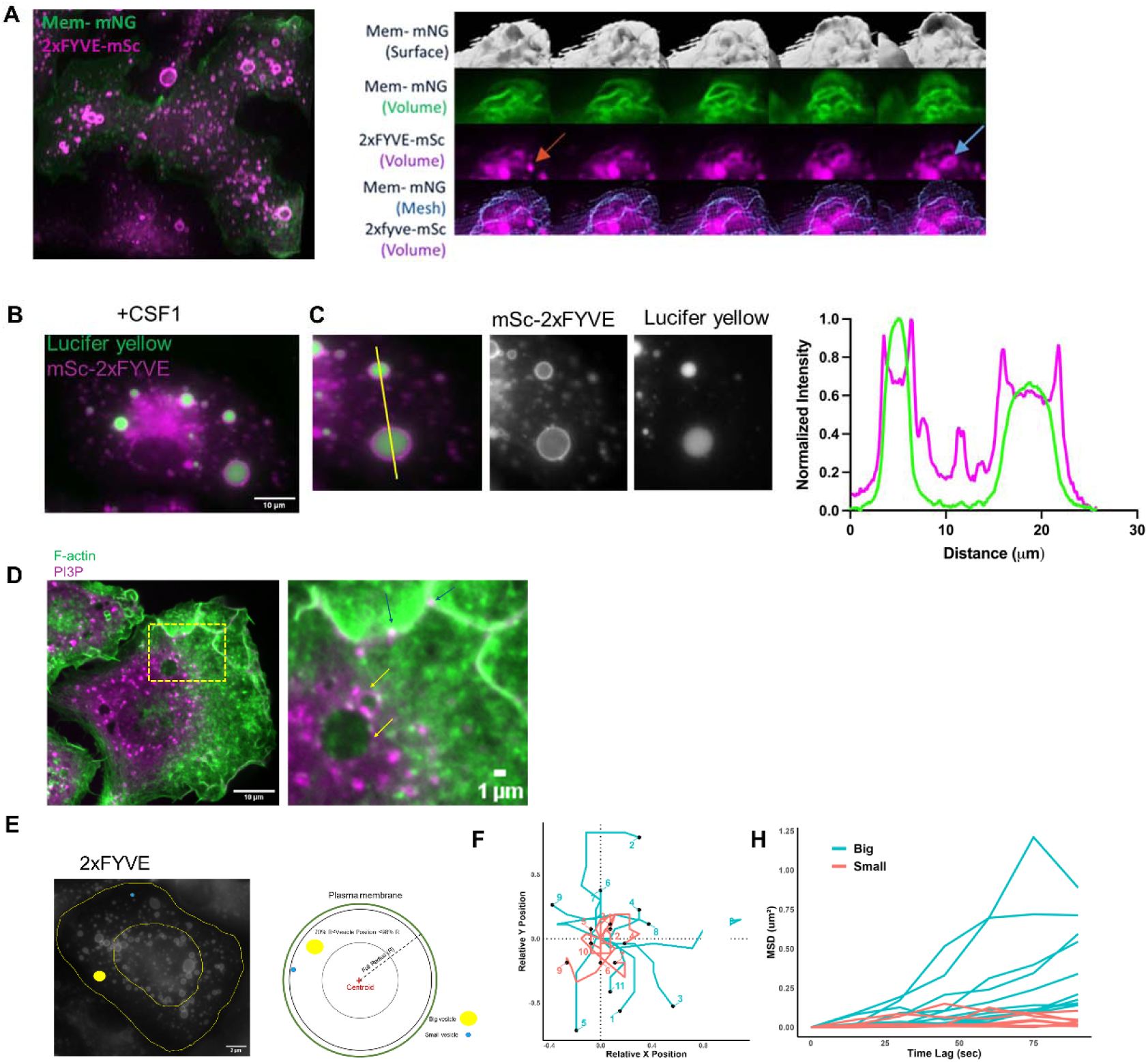
LLS imaging of shows two distinct classes of PI3P positive vesicles. A) LLS image showing volumetric intensities of membrane (green) and 2xFYVE (magenta). B) Confocal imaging of WT BMDMs expressing 2xFYVE-mSc (magenta) stimulated with CSF1 in the presence of Lucifer yellow dye (green) for 2.5 min, washed and imaged. Representative individual XY optical sections show the localization Lucifer yellow dye inside 2xFYVE positive vesicle in macropinosomes. Scale bar = 10µm. C) Intensity line-scan analysis of the XY optical section was performed along the yellow line indicated across the vesicle. Relative fluorescence intensities of 2xFYVE-mSc (magenta) and Lucifer yellow (green) demonstrate their co-localization and recruitment at open macropinosomes. D) Confocal imaging of WT BMDMS cultured on coverslips fixed and permeabilized as described in the methods section. Cells were starved overnight and stimulated with CSF1 for 5 minutes. Cells were stained with PI3P-snap-Alexa-647 and Phallodin. Brightness settings are equal across all images. Scale bar = 10µm. E) Schematic of the workflow used to quantify 2xFYVE-positive vesicle motility during macropinocytosis. WT BMDMs expressing 2xFYVE-mSc were stimulated with CSF1 and subjected to live-cell volumetric imaging, followed by particle detection from z-stack projections and reconstruction of vesicle trajectories. Individual vesicles were tracked in two dimensions and analyzed relative to the cell centroid. Scale bar=2 µm. F) Trajectory analysis of 2xFYVE-positive vesicles revealed heterogeneous motility behaviors following CSF1 stimulation. G) Mean square displacement (MSD) analysis of vesicle tracked in Figure F. This further demonstrated the coexistence of directed and confined movement modes within the vesicle population, consistent with dynamic remodeling during macropinosome maturation.

To quantify vesicle dynamics, we developed a tracking pipeline using epifluorescence live-cell imaging of CSF1-stimulated BMDMs expressing 2xFYVE-mScarlet. Vesicles were identified from z-stack image series, manually segmented in Fiji/ImageJ, and their centroids used to reconstruct two-dimensional trajectories over time (Figure 5D-E). This analysis revealed substantial heterogeneity in vesicle motility: a subset of vesicles remained confined near the initial centroid, while others underwent pronounced intracellular displacement over the imaging period (Figure 5F). Mean square displacement (MSD) analysis distinguished two motility regimes (Figure 5H): vesicles with higher MSD (teal) displayed active, long-range transport, whereas those with lower MSD (red) exhibited confined, largely Brownian-like motion, consistent with the two morphological classes identified by LLS imaging. Similarly, while expressing 2xFYVE-mScarlet in WT, *Uvrag*^sgRNA^ and *Atg14*^gRNA^ BMDMs we found the 2xFYVE positive puncta were fewer in the *Uvrag*^sgRNA^cells, even when a cell was stimulated with CSF1, while a higher amount of 2xFYVE positive puncta was observed in the *Atg14^s^*^gRNA^ cell (Supplemental Figure S4).

### UVRAG and ATG14L differentially regulate macropinocytic activity and autophagic flux

To understand whether there is any relationship between macropinocytosis and autophagy, we investigated autophagic flux in both *Uvrag*^sgRNA^ and *Atg14*^sgRNA^ BMDM (Figure 6A, 6B). As expected, ATG14L deficiency impaired autophagy, as indicated by accumulation of p62. However, we found no effect of UVRAG depletion on p62 accumulation. Of note, these cells are cultured in full growth medium and do not lack any nutrients. Interestingly, the relatively low p62 levels in WT BMDM indicate a constitutive autophagy program, that is impaired in the absence of ATG14. These results suggest that there is no direct one-one correlation of autophagy and macropinocytosis. Whether the increase in macropinocytosis in ATG14, is triggered by a metabolic demand or due to the increased activity of VPS34-II remains to be determined.

**Figure 6.**
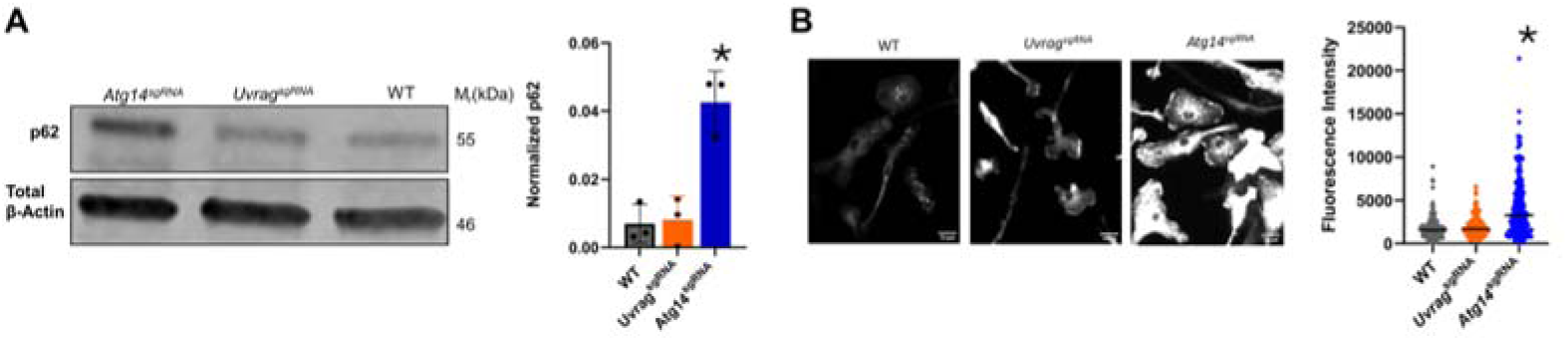
UVRAG and ATG14L differentially regulate macropinocytic activity and autophagic flux. A) Representative western blot showing p62 in the cells Quantification of p62. Bar graphs represent the mean ± SD of three independent experiments. *p < 0.05. B) BMDM cells were grown in BMM for 24 hours and stained primary antibody p62 overnight in 4 C and stained with secondary antibody. Brightness settings are equal across all images. Scale bar = 10 µm

## Discussion

Macropinocytosis depends on coordinated phosphoinositide signaling to regulate membrane remodeling, cup formation, membrane fusion and vesicle maturation. Although class I phosphoinositide 3-kinases (PI3Ks) and their products are well-established orchestrators of membrane ruffling and macropinocytic cup formation (Araki et al. 1996; Quinn et al. 2021; Welliver & Swanson 2012), the precise contributions of Class III PI3K signaling during macropinosome have remained unclear. The prevailing view has historically restricted PI3P function to post-scission endosomal maturation, where Rab5 recruits Class III PI3K to nucleate early endosomal machinery after the nascent vesicle has sealed (Christoforidis et al. 1999; Feliciano et al. 2011). Here, using CRISPR/Cas9 whole-genome screens in BMDMs we identified PI(3)P, synthesized by VPS34-II, acting directly at the plasma membrane prior to plasma membrane fusion and macropinocytic cup sealing.

Our findings challenge the view that early cup-associated PI(3)P is generated via sequential PIP_3_ dephosphorylation (Welliver & Swanson 2012; Yoshida et al. 2015). Selective pharmacological blockade of class I PI3Ks eliminated - recruitment without disrupting PI3P formation. Conversely, selective inhibition or genetic disruption of *Uvrag* abolished cup-associated PI(3)P without impairing Class I PI3K-dependent phosphorylation. Or membrane ruffling. Our live-cell imaging and membrane-impermeant FM4-64 labeling revealed that PI3P accumulates directly within open, unsealed macropinocytic cups prior to scission. Furthermore, time-lapse imaging confirmed that PI(3)P recruitment is mandatory for macropinsome formation as PIP_3_ positive cups that failed to acquire PI3P aborted and collapsed back into the plasma membrane. This failure to seal is similar to the observations in HT1080 cells, where SAR405-mediated VPS34 inhibition caused Phafin2-mNeonGreen positive nascent macropinosomes to regress and re-fuse with the plasma membrane (Spangenberg et al. 2021). These data demonstrate that class I PI3Ks and VPS34-II operate in parallel, non-redundant signaling pathways.

The reciprocal effects of UVRAG and ATG14 were among the most striking findings of our screen. VPS34 complexes I and II share a common VPS34–VPS15–Beclin1 core but contain mutually exclusive ATG14 or UVRAG subunits (Backer 2016; Li et al. 2012; Ohashi et al. 2019). Thus, loss of ATG14 may increase the assembly of UVRAG-containing VPS34 complex II, enhancing localized PI(3)P production. Consistent with this, ATG14-depleted cells showed increased PI(3)P on nascent macropinosomes whereas UVRAG depleted cells showed the opposite phenotype. These results suggest that spatially localized VPS34 activity, rather than total PI(3)P abundance, regulates macropinocytosis. Importantly, UVRAG loss did not cause p62 accumulation, indicating that its role in macropinocytosis is independent of defective autophagy. Together, our findings support a model in which UVRAG-containing VPS34 complex II promotes localized PI(3)P production during macropinocytosis, while ATG14 may negatively influence this process by regulating the distribution of the shared VPS34 core.

Beyond closure, PI(3)P may also support recycling of membrane and fusogenic machinery needed for repeated rounds of macropinocytosis. Our LLSM data are consistent with this as we observed distinct PI(3)P-positive populations including large nascent macropinosomes and as well as smaller, more dynamic puncta suggesting the presence of functionally separable compartments involved in closure versus recycling. Known PI(3)P effectors, including the BEACH-domain containing *Wdfy3/*ALFY (Simonsen et al. 2004), Plekhf2/PHAFIN2 (Schink et al. 2021), and PX-BAR domain sorting nexins (Kerr et al. 2006; Lim et al. 2008) were hits in our screen which offer plausible mechanisms linking VPS34-generated PI(3)P to membrane remodeling and tubulation. We further speculate that UVRAG-containing VPS34 complex II may coordinate with the HOPS tethering complex to support post-closure trafficking, consistent with reports that UVRAG physically and functionally cooperates with HOPS to direct endolysosomal trafficking (Bo et al. 2020), potentially linking PI(3)P-dependent macropinosome maturation to membrane recycling and lysosomal fusion.

## Materials and Methods

### Mice

Transgenic C57BL/6J mice constitutively expressing Cas9 from the H11 locus were obtained from the Jackson Laboratory (Strain# 028239). All experimental animals were maintained under pathogen-free conditions. Mice were humanely euthanized via carbon dioxide CO_2_ inhalation followed by cervical dislocation in strict accordance

### Primary cell cultures and cell lines

Bone marrow-derived macrophages (BMDMs) were harvested from the femurs of 8-to-12-week-old Cas9-expressing C57BL/6J mice. Mice were euthanized using CO2 inhalation and cervical dislocation, femurs were dissected from the mice, and bone marrow was harvested by flushing Dulbecco’s phosphate buffer saline (DPBS) through the bone using a needle and syringe ^3^. Cells were washed and cultured on non-tissue culture-treated plates in 5% CO2 and 37°C in bone marrow medium (BMM) consisting of DMEM, 20% heat-inactivated FBS, 50 ng/ml recombinant murine CSF1, 10,000 IU penicillin, 10 mg/ml streptomycin, and 0.0004% 2-mercaptoethanol. On day four post-isolation, non-adherent cells were discarded, and the adherent cells were cultured as macrophages in BMM. BMDMs were transduced on day 4 or 5 post isolation.

Human embryonic kidney HEK293T cells were maintained in DMEM supplemented with 10% HI-FBS and 100 U/mL penicillin-streptomycin.

### Key resources and reagents

sgRNA sequences used for targeted gene disruption (Supplementary Tabel S3), primers for genotyping (Supplementary Tabel S4), antibodies (Supplementary Table S5), cell culture medium (Supplementary Table S6), drugs and cytokines (Supplementary Table S7), commercial assays (Supplementary Table S8), plasmids (Supplementary Table 9), Other Reagents and Supplies (Supplementary Table S10). Mouse Brie CRISPR knockout pooled library was a gift from David Root and John Doench (Addgene #73633). PsPAX2 was a gift from Didier Trono (Addgene plasmid # 12260; http://n2t.net/addgene:12260; RRID: Addgene_12260). pCMV-VSV-G was a gift from Bob Weinberg (Addgene plasmid #8454; http://n2t.net/addgene:8454; RRID:Addgene_8454).

### Lentiviral production and transduction

Brie library amplification, lentiviral production, and titer calculations were performed as described(Joung et al. 2017) with minor modifications. Briefly, the Brie library plasmids were amplified in Stbl3 *Escherichia coli* (New England Biolabs, Ipswich, MA) and sequenced to assess sgRNA representation. To produce lentiviral particles for transduction, HEK293T cells were seeded onto 10-cm plates in 10% FBS plus DMEM before transfection with 6 µg sgRNA LentiGuide-Puro plasmid, 6 µg psPax2 plasmid, 1 µg pVSVG plasmid(STEWART et al. 2003), and 24 µg polyethyleneimine for 24 h. Lentiviral supernatant was harvested after 48 h and stored at −80°C. To calculate the functional viral titer, macrophages were transduced with increasing volumes of lentivirus in BMDM plus 1 µM cyclosporin A. After 48 h, the transduction solution was replaced with BMDM plus 5 µg/ml puromycin for selection. After selection, the surviving cells were counted to determine the percentage of transduced cells relative to nonselected control wells. For each screen, we transduced ∼ 5 × 10^7^ cells with a viral titer to achieve at least 50% cell death following antibiotic selection. The low titer was to ensure that the majority of transfected cells contained only one guide per cell. Cells were cultured for at least 8 days following puromycin selection to allow for gene disruption and protein turnover. Similar procedures were followed for lentiviral delivery of sgRNA for targeted single gene disruptions.

### Whole genome screen workflow

For the whole-genome screen Lucifer yellow uptake assays, cells were starved overnight in CSF1-free medium (DMEM plus 10% FBS) and subsequently exposed to 500 mg/ml Lucifer yellow for 30 min in the presence of 200 ng/ml CSF1 and PMA in DMEM plus 10% FBS preequilibrated to pH 7.4 (Figure 1). Of note, BMDM internalize CSF1R from the surface of the cell in basic medium(Huang et al. 2024). After 30 min, the cells were gently washed with warm PBS, incubated on ice for 5 min in calcium- and magnesium-free PBS, and pipetted to facilitate detachment. Detached cells were sorted with a BD FACS Jazz flow cytometer based on Lucifer yellow fluorescence with the highest (high Lucifer yellow uptake) and lowest (low Lucifer yellow uptake) quintiles sorted for sequencing of the sgRNA inserts. Approximately 10% of the cells were collected before sorting to assess sgRNA insert distribution in the cell population (Presort). After sorting, the cells were pelleted and stored at −80°C before DNA purification. In the targeted knockout studies, Lucifer yellow uptake assays were conducted as indicated in the figure legends using Cytoflex and quantified using FlowJo^TM^10.

### PCR and next-generation sequencing of gRNA inserts

Genomic DNA was extracted from pellets with the GeneJET genomic DNA extraction kit and the sgRNA library was amplified for sequencing as described(Joung et al. 2017). Briefly, genomic DNA was amplified equally in each reaction with an equimolar mix of P5 staggered primers and a unique P7 primer (Supplemental Table S4) containing an index sequence for sample identification with a Phusion polymerase kit. Following genomic DNA amplification, the PCR product was pooled and the 357 bp product was gel purified with the wizard gel and PCR clean up kit for sequencing with the Illumina Nextseq 500 system high-output kit with 75 bp read length. Sequencing quality was assessed within the Illumina dashboard and read count summaries and mapping data from MAGeCK (Supplemental Figures S4 and S5). Cut adapt, version 1.18 *(Martin, 2011) was used to remove sequencer adapter regions to produce FASTQ files containing the 20 bp sgRNA sequences.

### Statistical analysis of sequencing data and gene ranking

Model-based analysis of genome-wide CRISPR-Cas9 Knockouts (MAGeCK), version 0.5.9 (Li et al. 2014) identified and ranked enriched sgRNAs in the highest and lowest quintiles based on dextran uptake. The “—count” function mapped sgRNAs sequences processed with Cutadapt to the Brie library sequences file and generated read counts for each sgRNA. The “-test” command was used to compare sgRNA read counts for each sgRNA in the low- and high-fluorescence populations and rank each sgRNA for enrichment to identify sgRNAs, and therefore genes, that regulate dextran uptake. The “-control-sgRNA” command was used with the “-test” command with identify 1000 control noncoding sgRNAs used as a control for read count normalization. (Ge et al. 2019; Kanehisa et al. 2020; Luo & Brouwer 2013)

### Cut-site sequencing to confirm gene disruption

Genome editing efficiency was verified by targeted amplicon sequencing of CRISPR cut sites. Between 0.5 × 10^6 and 2 × 10^6 puromycin-selected BMDMs were harvested 7–10 days after lentiviral transduction and pelleted by centrifugation. Cell pellets were washed with PBS, frozen at −80°C, and genomic DNA was isolated using the GeneJET Genomic DNA Purification Kit. PCR primers flanking each sgRNA target site were designed using CHOPCHOP software with an expected amplicon size of 350–450 bp. Target loci were amplified using Phire Hot Start II DNA Polymerase in 50-μL reactions containing 0.3 μM forward and reverse primers and up to 1 μg genomic DNA. PCR products were purified using AMPure XP magnetic beads, quantified, and submitted to Plasmidsaurus for nanopore sequencing using the Premium PCR sequencing service. Sequencing data were analyzed using CRISPResso2(Clement et al. 2019) to determine insertion/deletion frequencies and the proportion of remaining wild-type alleles. Editing efficiencies were calculated from the percentage of modified alleles relative to total aligned sequencing reads. Gene-specific primer sequences are provided in Table S4.

### Macropinocytosis assays

Macropinocytosis was quantified by measuring the uptake of the fluid-phase tracer Lucifer Yellow. Bone marrow-derived macrophages (BMDMs) were serum-starved overnight in CSF1-free medium consisting of DMEM supplemented with 10% fetal bovine serum (FBS). Cells were stimulated with 200 ng/mL recombinant mouse CSF1 in the presence of 500 μg/mL Lucifer yellow. Stimulation times are indicated in the corresponding figure legends. For flow cytometric analyses, cells were detached by incubation in calcium- and magnesium-free PBS on ice for 5 min followed by gentle pipetting. Cellular Lucifer yellow fluorescence was quantified using a CytoFLEX flow cytometer and analyzed using FlowJo v10 software. For microscopy-based experiments, at the completion of each stimulation, cells were rapidly washed with in Hanks’ Balanced Salt Solution (HBSS) and imaged in using an Olympus IX83 inverted fluorescence microscope maintained at 37°C. Fluorescence excitation was provided by an X-Cite Turbo illumination system, and emitted fluorescence was collected using a Quad OSF-QUADPLEDBX3 filter cube. Images were acquired using either a 40× air objective (NA 0.95) or a 60× oil immersion objective (NA 1.42) with a cooled CCD camera. Quantification of Lucifer Yellow fluorescence was performed using Fiji/ImageJ as described in the corresponding figure legends.

#### Pulse-Chase Experiments

BMDMs were serum-starved overnight in CSF1-free medium before stimulation with 200 ng/mL recombinant mouse CSF1 in the presence of 500 μg/mL Lucifer Yellow for the pulse period indicated in the figure legends. Following the pulse, cells were washed extensively with phosphate-buffered saline (PBS) to remove extracellular dye and incubated in fresh DMEM at 37°C for the indicated chase period. At each chase endpoint, cells were washed with PBS, transferred to ice, detached in calcium- and magnesium-free PBS, and analyzed by flow cytometry. Lucifer Yellow fluorescence remaining within each cell was quantified using a CytoFLEX flow cytometer and analyzed with FlowJo v10 software.

### Western Blot

Drug treatment and CSF-1 stimulation were performed as described in the corresponding figure legends. Following treatment, cells were lysed in Mammalian Protein Extraction Reagent (M-PER) supplemented with Halt protease inhibitor cocktail and phosphatase inhibitor cocktail. Cell lysates were incubated on ice for 10 min and lysates were cleared by centrifugation at 18,000 × g for 15 min at 4°C. Protein concentration was determined using the Pierce BCA Protein Assay Kit according to the manufacturer’s instructions. Equal amounts of protein (20 µg) were loaded per well on 4–20% SDS–PAGE precast gels and separated by electrophoresis. Proteins were then transferred to polyvinylidene fluoride (PVDF) membranes by wet transfer in transfer buffer containing 20% methanol. Membranes were blocked with 5% non-fat dry milk in bovine serum albumin and incubated with primary antibodies diluted according to the manufacturer’s recommendations overnight at 4 °C. Following incubation with appropriate secondary antibodies, membranes were imaged using the LI-COR Odyssey Fc imaging system. Band intensities were quantified using Image Studio Lite software.

### Construction of the membrane and 2xFYVE probes

The membrane probe was constructed by combining the membrane localization motif (MGCVCSSNPE) from Lck in frame with mNeonGreen in the pLJM1 backbone containing the puromycin resistance gene. Mscarelet-2XFYVE was constructed as follows: 2xFYVE sequence from(Gillooly et al. 2000; Quinn et al. 2021) with one FYVE domain codon optimized to mouse, was synthesized and inserted into a pLenti plasmid containing mScarlet-I the blasticidin resistance cassette. FVYE sgRNA sequence: TCCGAAAGTGATGCCATGTTCGCTGCTGAAAGAGCCCCTGACTGGGTGGATGCTGAGGAATGCCATCGGTGCAGAGTACAGTTTGGGGTGGTGACCCGCAAGCATCACTGCCGAGCATGTGGGCAGATCTTCTGTGGCAAGTGCTCCTCCAAGTACTCCACCATCCCCAAGTTCGGCATTGAGAAGGAGGTGCGCGTGTGTGAGCCCTGCTATGAGCAGCTGAACAAGAAGGCA

### Visualization of AktPH, 2xFYVE and UVRAG dynamics during macropinocytosis

Day 4 BMDMs were transduced with lentiviral vectors encoding pTwist Lenti SFFV-Uvrag-mScarlet-I, pLJM1-mScarlet-2×FYVE, pLJM1-AktPH-mScarlet, or pLJM1-Lck-mNeonGreen. Cells were maintained in culture until days 7–8 before live-cell imaging experiments.

For Lucifer Yellow uptake experiments, transduced macrophages were plated onto 35-mm glass-bottom imaging dishes and serum-starved overnight in CSF1-free medium. Cells were stimulated with CSF1 in the presence of Lucifer Yellow, washed with DPBS, and immediately imaged by confocal microscopy.

To distinguish sealed macropinosomes from open circular ruffles, serum-starved macrophages were stimulated with 200 ng/mL CSF1 for 1 min at 37°C. Cells were rapidly washed with ice-cold PBS and incubated with 20 μM FM4-64 for 1 min on ice to label the plasma membrane. Samples were immediately transferred to the microscope for image acquisition. Sealed macropinosomes were identified as intracellular Lucifer yellow-positive structures lacking FM4-64 labeling, whereas open membrane ruffles remained continuous with the plasma membrane and retained FM4-64 fluorescence.

For inhibitor studies, macrophages were pretreated with pharmacological inhibitors for 10 min before CSF1 stimulation. Cells were maintained in the continued presence of inhibitor throughout imaging. Time-lapse imaging was performed using an Olympus live-cell imaging system for 5 min immediately following stimulation under temperature-controlled conditions.

Representative images shown in the figures were processed identically for brightness and contrast using Fiji/ImageJ. Quantitative image analysis was performed on raw image files using identical thresholding parameters for all experimental groups.

### Immunofluorescence staining

Bone marrow-derived macrophages (BMDMs) were seeded onto ethanol-flamed 12-mm glass coverslips and cultured overnight. Following stimulation, cells were fixed in 4% paraformaldehyde containing 0.2% glutaraldehyde in phosphate-buffered saline (PBS) for 12 min at room temperature. Cells were permeabilized with 0.3% Triton X-100 for 1 h and subsequently blocked with 5% bovine serum albumin (BSA) in PBS for 1 h at room temperature. Primary antibodies were diluted in 1% BSA in PBS and incubated with samples overnight at 4°C. After washing with PBS, cells were incubated with the appropriate fluorescently conjugated secondary antibodies for 1 h at room temperature. Nuclei were counterstained with DAPI-containing Fluoromount mounting medium.Wide-field fluorescence images were acquired using an Olympus IX83 inverted microscope equipped with either a 40× air objective (NA 0.95) or a 60× oil immersion objective (NA 1.42). Confocal imaging was performed using a TILL Photonics Andromeda spinning-disk confocal microscope equipped with a 60× oil immersion objective. Identical acquisition settings were maintained within each experiment to enable quantitative comparison among treatment groups.

### Lattice light sheet microscopy

Macrophages were prepared for LLS imaging 24 h prior to imaging using 5 mm glass coverslips. The coverslips were soaked in 90–100% ethanol and flame cleaned using a butane flame. Approximately 5 flame cleaned coverslips were placed per well of a 12-well plate each containing 1 ml of culture media. Cells were added to each well during the cell culture process at ∼3 × 10^5^ cells in each 3.5 cm^2^ (12-well plate) for imaging. The FLMs were incubated on the flame-cleaned glass coverslips in culturing media for 24 h prior to imaging. The coverslips were transferred to the LLSM bath that was filled with 7 ml of Leibovitz’s L-15 Media (supplemented with 1.7 mM glucose) at ∼37 °C.

The LLSM is a replica of the design described by Chen et al.25, built under license from HHMI. Volumetric image stacks were generated using dithered square virtual lattices (Outer NA 0.55, Inner NA 0.50, approximately 30 µm long) and stage scanning with 0.5 µm step sizes, resulting in 254 nm deskewed z-steps. Excitation laser powers used were 18 µW (488 nm) and 22 µW (561 nm), measured at the back aperture of the excitation objective. The emission filter cube (DFM1, Thorlabs) comprised a quadband notch filter NF03-405/488/561/635 (Semrock), longpass dichroic mirror Di02-R561 (Semrock), shortpass filter 550SP (Omega) on the reflected path, and longpass filter BLP01-561R (Semrock) on the transmitted path; the resulting fluorescence was imaged onto a pair of ORCA-Flash4.0 v2 sCMOS cameras (Hamamatsu). The camera on the reflected image path was mounted on a manual x–y–z translation stage (Newport 462-XYZ stage, Thorlabs), and the images were registered using 0.1 µm Fluoresbrite YG microspheres (Polysciences). The image capturing rates varied between 5 and 10 s per volume using 8–12 ms planar exposures depending on the brightness of the cell and imaging region.

### Live cell imaging and Vessicle tracking

Day 4 BMDMs were transduced with lentiviral contructs encoding pLJM1-mScarlet-2xFYVE using lentiviral transduction system. Cells were cultured until day 7-8 before imaging experiments.

Cells were stimulated with CSF1 and imaged using an Olympus epifluorescence microscope. Time-lapse images were acquired at 15 s intervals. Vesicle tracking was performed on seven z-stack projections. To preferentially analyze peripheral vesicles, structures located between 75– 98% of the radial distance from the cell centroid were selected for analysis. Individual vesicles were manually segmented in Fiji/ImageJ by drawing regions of interest (ROIs) for each time point. Vesicle centroid coordinates and Feret diameters were subsequently extracted for downstream quantitative analysis.

Trajectory reconstruction and motility analysis: Two-dimensional vesicle trajectories were reconstructed from centroid coordinates over time. For comparative visualization of vesicle displacement, trajectories were normalized by translating the initial position of each vesicle to a common origin (0,0). These origin-centered trajectories (“rose plots”) were used to assess directional persistence and displacement patterns across vesicle populations.

Quantitative analysis of vesicle motility was performed in R using the tidyverse and ggplot2 packages. Mean square displacement (MSD) was calculated for each vesicle across increasing time intervals (τ) according to:

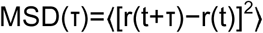

where r(t) represents vesicle position at time t, and images were acquired at 15 s intervals. To characterize vesicle transport behavior, MSD curves were fitted to a power-law model:

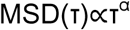

### Purification, labeling of PI(3)P biosensors and staining

The recombinant PI(3)P biosensor comprised of 2xFYVE domain of HRS and SNAP tag was obtained from Addgene ( #211508 ) and expressed in Escherichia coli with minor modifications to the published protocol (Maib et al. 2024). Brifely, the recombinant PI(3)P biosensor were expressed in Rosetta^TM^ 2(DE3) Competent cells (71397, Sigma-Aldrich) BL21 bacterial cells using standard approaches. Transfected bacteria were grown in LB containing antibiotic at 37°C to OD600 = 0.8. The bacteria was induced with 1 mM ITPG at and grown at 18°C overnight. Cells were pelleted, lysed using B-Per^TM^ (78266, thermos scientific) and centrifuged. Cleared lysates were passed through a 0.45-µM filter (Sartorius), and protein was purified by Ni2+ affinity chromatography using 5 ml His-Trap HP column (Cytiva) against an increasing gradient of standard buffer containing 250 mM Imidazole using Syringe pump. Desalting of the sample was done by using Zeba spin Desalting Columns (Thermofisher Scientific). All proteins were aliquoted, frozen in liquid nitrogen, and stored at −80°C. Aliquots were labeled with SNAP-Surface Alexa Fluor 647 (NEB) in a 2:1 M excess for 2–3 h on ice in a standard buffer. Excess dye was removed by dialysis Cassettes (PI66380, Thermo Scientific) against standard buffer at 4°C overnight.

A total of 20,000 cells were grown in 96 well plate in CSF1-free medium (DMEM plus 10% FBS) overnight. Cells were stimulated with 200 ng/ml CSF1 for the time indicated in the figure legends. The culture medium was removed, and cells were fixed using pre-warmed 4% paraformaldehyde and 0.2% glutaraldehyde for 20 minutes at room temperature. Residual aldehydes were quenched by two rapid NH_4_Cl washes, followed by incubation with 50 mM NH_4_Cl in PBS for 20 minutes. Cells were stained, blocked, and permeabilized at the same time in 5% BSA + 0.5% (vol/wt) Saponin (558255, Millipore) in PIPES (J63617.AE,Thermofisher) with the addition of labeled biosensors to a final concentration of 500 nM for 45–60 min on ice. For staining, a standard metal heat block was inverted and cooled down in an ice bucket. Cell were washed three times with ice-cold PIPES and post-fixed with 2% PFA in PBS on ice for 10 min before being returned to room temperature for another 10 min. Cells were washed with 50 mM NH_4_Cl in PBS at room temperature. Nuclei were stained with DAPI and. phallodin. Images were acquired on a wide-field epifluorescence Olympus IX83 microscope using 40x air objective and 60x oil objective. Confocal images were acquired using a TIL photonics Andromeda Spinning Disk Confocal (TILL Photonics, Munich, Germany) with a 60x oil objective.

### Surface marker immunostaining

A total of 1.5 × 10^5^ cells were suspended in 1% FBS plus 1.5 µg/ml Fc block in PBS for 15 min on ice according to the manufacturer’s instructions. After 15 min, Phycoerythrin (PE)-conjugated anti-MRC1 or isotype control (1:80) was added to each sample and incubated for 15 min on ice. The samples were then washed three times and fluorescence intensity measured with a BD Accuri flow cytometer. At least 10^4^ cells were analyzed for each experiment and live cells gated using forward and side scatter.

### Statistical analyses

Geometric means and medians for flow cytometry data were calculated using FlowJo software (version 10.6; BD Biosciences, Franklin Lakes, NJ). Western blot images were quantified using Image Studio Lite software (version 5.2; LI-COR Biosciences, Lincoln, NE). All statistical analyses were conducted using GraphPad Prism (version 8.0; San Diego, CA) and R (version 4.3.2) within RStudio. Data are presented as mean ± 95% CI. Differences between two groups were assessed using Student’s t-test. Comparisons among three or more groups were evaluated using one-way ANOVA followed by Dunnett’s post hoc test. Significance was set at p < 0.05. Sample sizes and specific p-values are indicated in the respective figure legends.

