## Supplemental Figure for "PI(3)P Signaling by VPS34 Complex II Orchestrates Macropinocytosis"

**Supplemental Materials**

Supplementary Table S1. MAGeCK gene summaries

Supplementary Table S2. MAGeCK pair analysis

Supplementary Table S3. sgRNA sequences used for targeted gene disruption.

| Target | Source | gRNA Sequencing (5' to 3') | Brie |
|---|---|---|---|
| *Atg14* | Addgene (submission in transit) | CTGCGTGGCGTAGCCCAGCG | Brie_078539 |
| *Beclin* | Addgene (submission in transit) | *GGATGACGAACTCAAGAGTG* | Brie_024265 |
| Non-Targeting gRNA | Addgene (submission in transit) | AAAAAGTCCGCGATTACGTC | Brie_078638 |
| *Pik3c3* | Addgene (submission in transit) | AGCCTGTAAGAACTCAACAC | Brie_056572 |
| *Pik3r4* | Addgene (submission in transit) | GGGTACTGATGCGATCGTAG | Brie_042011 |
| *Sh3glb1* | Addgene (submission in transit) | *GATTACCCGACTCCTGCTTG* | Brie_023815 |
| *Uvrag* | Addgene (submission in transit) | CAGCGATGTTCCGGAGATGT | Brie_044271 |

Supplementary Table S4. Primers for Genotyping

| Target | Forward (5' to 3') | Reverse (5' to 3') |
|---|---|---|
| *Atg14* | TAGATGTATACTGTGGCTGCGG | GCTCTCAGCAAAGGTTTGAGAT |
| *Beclin* | AGCACCTGGAATTTGAGACATT | CTTCTATGCAGAGGTGTGTTGC |
| *Pik3c3* | CCTCTTAGGCAACCTCAGAAAA | TTCAAAGGGGTGATACAAAACC |
| *Pik3r4* | AACTGAAAATCAGGCTCCACTC | GGGAGCTATGTAGCATGTCCTC |
| *Sh3glb1* | TTGTGTCAGTTGTGTTTGGTGA | GAGAGGTAAATGCCACAGGAAG |
| *Uvrag* | CTGCTGTTTGAGTGAGATGAGG | AGTAGAAGAAGGCACTGCAAGC |

Supplementary Table S5. Antibodies

| Reagents | Source | Identifier | Dilution and working concentration |
|---|---|---|---|
| Akt (pan) (40D4) Mouse Monoclonal Antibody, Clone | Cell signaling technology | 2920 | 1:500, 5% BSA |
| Alexa Fluor® 488 anti-mouse F4/80 Antibody, Clone: BM8 | Biolegend | 123119 | 5 ug/ml |
| Alexa Fluor® 488 Rat IgG2a, κ Isotype Ctrl Antibody, Clone: RTK2758 | Biolegend | 400525 | 5 ug/ml |
| Anti-mouse IgG (H+L) (DyLight® 680 Conjugate) | Cell signaling technology | 5470 | 1:3000, 5% BSA |
| APC anti-mouse CD206 (MMR) Antibody /Mrc1, Clone: C068C2 | Biolegend | 141707 | 5 ug/ml |
| APC Rat IgG2a, κ Isotype Ctrl Antibody, Clone: RTK2758 | Biolegend | 400511 | 5 ug/ml |
| APC Rat IgG2b, κ Isotype Ctrl Antibody, Clone: RTK4530 | Biolegend | 400611 | 5 ug/ml |
| BD Pharmingen™ Alexa Fluor™ 488 Rat Anti-CD11b, Clone: M1/70 | Biolegend | 101211 | 5 ug/ml |
| beta-Actin (8H10D10) Mouse Monoclonal Antibody | Cell signaling technology | 3700 | 1:500, 5% BSA |
| Goat anti-Rabbit IgG (H+L) Secondary Antibody, DyLight™ 800 4X PEG | Invitrogen, Thermofisher Scientific | SA5-35571 | 1:3000, 5% BSA |
| PE anti-mouse CD115 (CSF-1R) Antibody, Clone:AFS98 | Biolegend | 135505 | 5 ug/ml |
| PE Rat IgG2a, κ Isotype Ctrl Antibody, Clone: RTK2758 | Biolegend | 400507 | 5 ug/ml |
| Phosphatidylinositol 3,4,5-trisphosphate Monoclonal Antibody, Clone:RC6F8 | Invitrogen, Thermofisher Scientific | A-21328 | 1:100, 2% BSA |
| Phospho-Akt (Ser473) (D9E) Rabbit Monoclonal Antibody | Cell signaling technology | 4060 | 1:500, 5% BSA |
| SQSTM1/p62 (D6M5X) Rabbit Monoclonal Antibody | Cell signaling technology | 23214 | 1:500, 5% BSA |

Supplementary Table S6. Cell Culture Medium

| Reagent | Identifier | Source |
|---|---|---|
| Dulbecco's Modified Eagle Medium (DMEM) | 30-2002 | ATCC |
| Heat-Inactivated Fetal Bovine Serum (HI-FBS) | S11150 | GeminiBio |

| | | |
|---|---|---|
| Penicillin/Streptomycin | MT30002CI | Corning |
| B mercapto ethanol | | Cayman Chemical |
| Recombinant Mouse M-CSF | 576404 | BioLegend |

Supplementary Table S7. Drugs and cytokines

| Reagent | Identifier | Source |
|---|---|---|
| A-66 | 17382 | Cayman chemical |
| Ampicillin | 14417 | Cayman chemical |
| AS-252424 | 10009052 | Cayman chemical |
| Blasticidin | 14499 | Cayman chemical |
| Chloramphenicol | 227920250 | Thermo Scientific |
| Cyclosporin A | 12088 | Cayman chemical |
| Halt Protease Inhibitor Cocktail | 78428 | Thermo Scientific |
| IC-87114 | 11589 | Cayman chemical |
| Kanamycin sulfate | 15321 | Cayman chemical |
| Ly294002 | 70920 | Cayman chemical |
| Mouse recombinant CSF1 | 576404 | BioLegend |
| Phorbol 12-myristate 13-acetate (PMA) | 10008014 | Cayman Chemical |
| Puromycin | 13884 | Fisher Scientific |
| SAR405 | 16979 | Cayman chemical |
| Vps34In-1 | 17392 | Cayman Chemical |

Supplementary Table S8. Commercial assays

| Reagent | Identifier | Source |
|---|---|---|
| AMPure XP Magnetic Beads | A63881 | Beckman Coulter |
| GeneJET Genomic DNA Purification Kit | K0721 | Thermo Fisher Scientific |
| Pierce BCA Protein Assay Kit | 23225 | Thermo Fisher Scientific |
| QIAgen midi prep kit | 27104 | Qiagen |
| Slide-A-Lyzer™ Dialysis Cassettes, 10K MWCO | PI66380 | Thermofisher |
| Zeba™ Spin Desalting Columns | 89890 | Thermo Fisher Scientific |

Supplementary Table 9. Plasmids

| Reagent | Identifier | Source |
|---|---|---|
| Mouse Brie CRISPR knockout pooled library | 73633 | David Root & John Doench / Addgene |
| pCMV-VSV-G | Plasmids# 8454 | Bob Weinberg / Addgene |

| | | |
|---|---|---|
| pLenti-mScarlet-I-2xFYVE | Addgene# [Pending] | This paper |
| pLJM1-AktPH-mScarlet | Addgene# [Pending] | This paper |
| pLJM1-mNeonGreen-Lck membrane probe | Addgene# [Pending] | This paper |
| psPAX2 | Plasmids# 12260 | Didier Trono / Addgene |
| pTwist Lenti SFFV-Uvrag-mScarlet-I | Addgene# [Pending] | This paper |
| pTwist Lenti SFFV-Uvrag- mcerulean3 | Addgene# [Pending] | This paper |
| Recombinant PI(3)P biosensor (2xFYVE-HRS-SNAP) | Plasmids# 211508 | Addgene |

Supplementary Table S10. Other Reagents and Supplies

| Reagent | Identifier | Source |
|---|---|---|
| 4% para formaldehyde | 158127 | Sigma aldrich |
| 10 x Tris/Glycine Buffer | 1610771 | Bio-Rad |
| Glutaraldehyde | 108790-50G | Millipore |
| Alexa Fluor 488 Phallodin | A12379 | Thermo Fisher Scientific |
| B-PER™ Bacterial Protein Extraction Reagent | 78266 | Thermo Fisher Scientific |
| Bond-Breaker™ TCEP Solution | 77720 | Thermofisher |
| Bovine Serum Albumin (BSA) | BP1600-100 | Fisher Scientific |
| Dulbecco's Phosphate Buffered Saline (calcium, magnesium) | SH3026401 | Fisher Scientific |
| GelCode Blue Safe protein | 1860957 | Thermo Fisher Scientific |
| FM464 | T13320 | Thermo Fisher Scientific |
| Fluromount-G with DAPI | 00-4959-52 | Thermo Fisher Scientific |
| Halt Protease Inhibitor Cocktail | 78428 | Thermo Fisher Scientific |
| Hank's Balanced Salt Solutions | MT21023CM | Fisher Scientific |
| HisTrap™ HP 5 mL Column | | Cytiva |
| IPTG | 15529019 | Invitrogen |
| LentiX- concentrator | | Takara Bio |
| Lucifer yellow | L453 | Thermo scientific |
| LB broth | BP9723 | Fisher Scientific |
| LB agar | BP9724 | Fisher Scientific |
| Mini protein gel TGX | 4561094 | BioRad |
| M-PER Mammalian Protein Extraction Reagent | 78503 | Thermo Scientific |
| HisPur Ni-NTA resin | 88221 | Fisher |
| NucBlue fixed cell DAPI | R37606 | Thermo Fisher Scientific |
| PIPES | 63617.AE | Thermo Fisher Scientific |

| | | |
|---|---|---|
| Saponin | 558255 | Millipore |
| mPAGE Color Protein Stand protein ladder | MPSTD4 | Millipore |
| SNAP-Surface® Alexa Fluor® 647 | S9136S | New England Biolabs |
| STBR safe DNA gel stain | S33102 | Fisher |
| Trypan blue | 25-900-CI | Corning |
| 0.05% Trypsin-EDTA (1X) | 25300-054 | Gibco |
| TGX-221 | 10007349 | Cayman chemical |
| Triton X-100 | CAS 9002-93-1 | Fisher Bioreagents |
| Mini-Protean TGX Gels | 64749368 | Bio-Rad |
| Methonal | A412-4 | Fisher |
| polyvinylidene fluoride (PVDF) membranes | 1620177 | BioRad |
| SNAP-Surface Alexa Fluor 647 (NEB) | S9136 | NEB |
| DPBS (1x) | 14040-133 | Gibco |
| Albumin Standard | 23209 | Thermo scientific |
| NEB 10-beta/Stable | B9035S | BioLabs |
| Mini Trans-Blot filter paper | 1703932 | Biorad |
| Immun-Blot PVDF Membranes for protein Blotting | 1620177 | BioRad |
| Phosphate buffer Saline (1x) -Ca/-Mg | SH3D256.02 | Cytiva |
| Empty FPLC Columns 1ml | MPPC001-8 | Biocomma |

### Supplemental figures

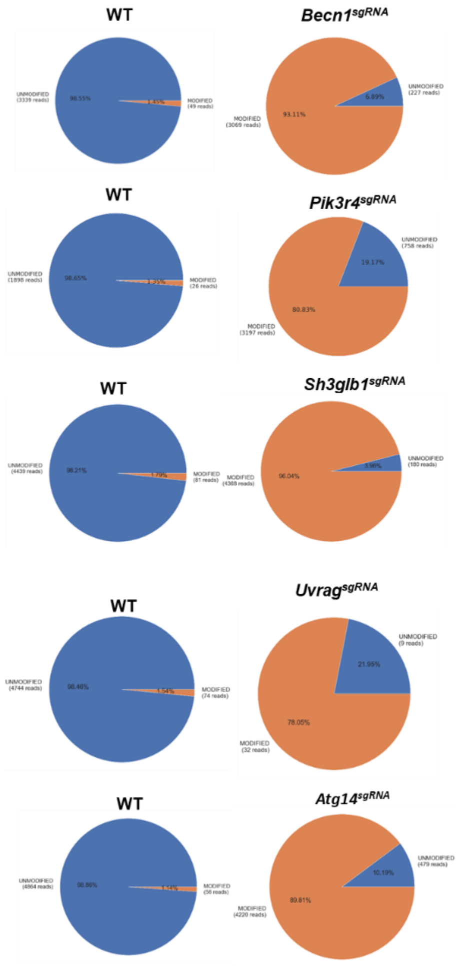


**Supplemental Figure S1.** Cut-site sequencing to confirm gene disruption. Genomic DNA was isolated from one million puromycin-selected macrophages 7 to 14 days post-gRNA transduction, amplified by PCR, and sequenced. Sequencing data was analyzed using CRISPResso2 to determine the percentage and number of reads corresponding to unmodified and modified alleles. The pie chart shows the percentage of modified and unmodified alleles, quantitatively assessing genome editing efficiency.

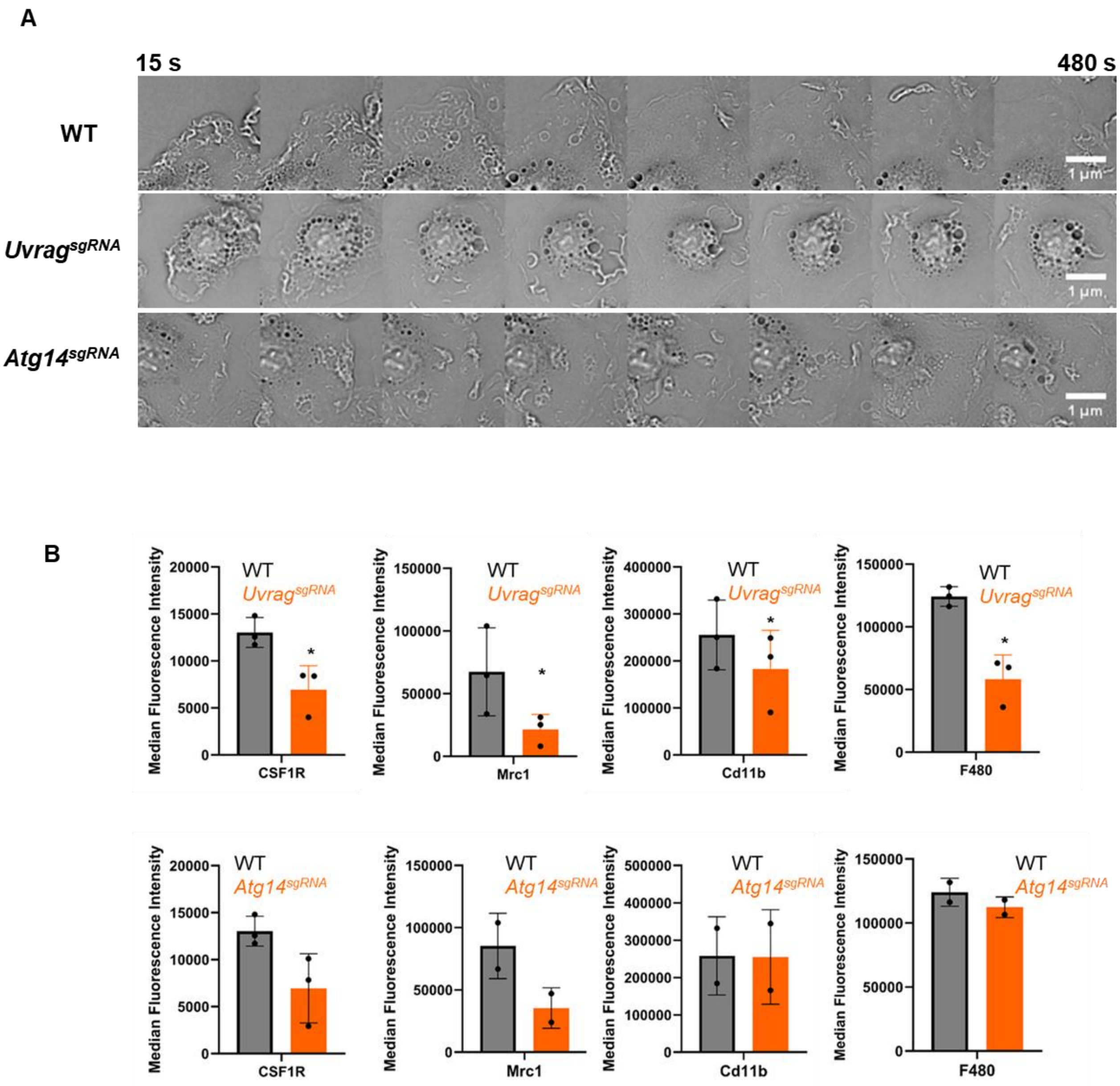


**Supplemental Figure S2.**

A) Time-lapse montage gallery of WT, *Uvrag*$^{sgRNA}$, and *Atg14*$^{sgRNA}$, BMDMs. Cells were starved overnight, stimulated with CSF1 for 8 min, and imaged in HBSS at 15-s intervals using an Olympus IX83 microscope with a 60× objective. Scale bar = 1 µm

B) BMDM with the indicated gene disruptions were immunostained for surface marker as described in method section and fluorescence intensity analyzed by flow cytometry. Bar graph shows the mean of replicate experiments, error bars are SD. * $p < 0.05$.

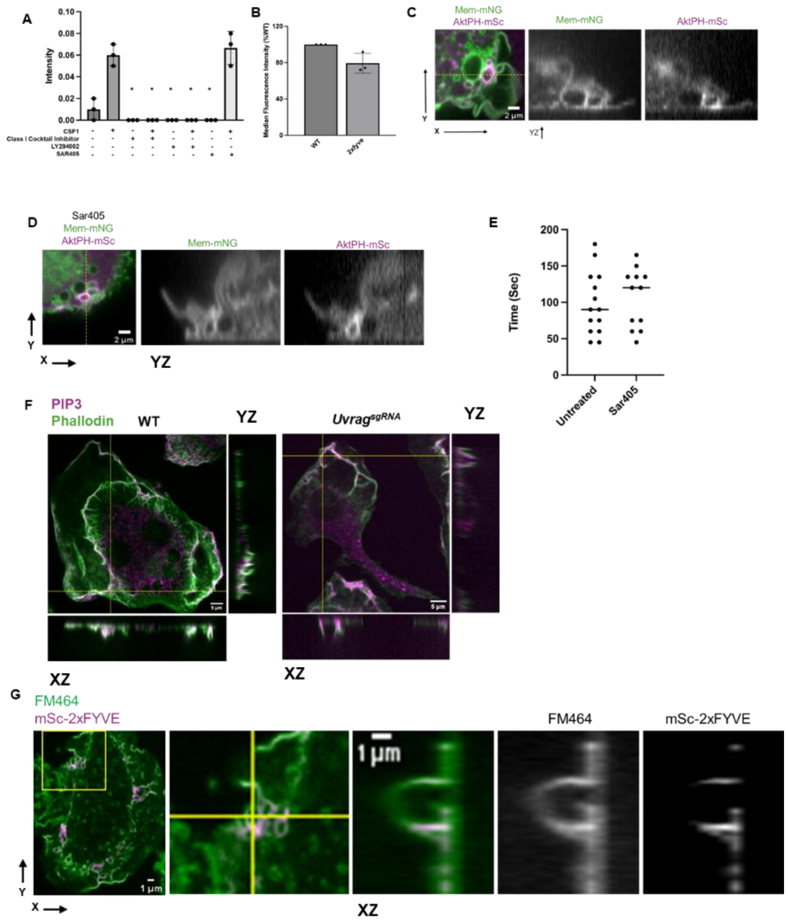


**Supplemental Figure S3**

A. Ratio of phosphorylated p-Akt to total AKT after PI3K inhibition depletion and CSF1 activation related to Figure 4A. (mean ± SEM; n = 3 replicates from 3 independent experiments).

B. Flow cytometry analysis of Lucifer yellow dye (405 nm) in WT with or without expressing 2xFYVE-mSc. BMDMs were starved overnight, stimulated with 200 ng/ml CSF1 for 30 minutes in the presence of 500 µg/ml Lucifer Yellow, washed and amount of dye uptake was measured by using flow cytometer. Median fluorescence intensity was normalized to WT BMDMs and corrected for autofluorescence using no-dye controls. Statistical analysis was performed using one-way ANOVA followed by Dunnett's multiple comparisons test (95% confidence interval, family-wise alpha = 0.05). Bar graphs represent the mean ± SD of three independent experiments. $^*p < 0.05$. Data was analyzed using FlowJO™ and GraphPad Prism.

C. Confocal imaging (related to Figure 4H) of BMDMs expressing Akt-PH-mScarlet (magenta) and Mem-mNG(green), stimulated with 200 ng/mL CSF1 for 2.5 min. Representative individual XY optical sections show the localization of Akt-PH-mScarlet (magenta) and Mem-mNG(green).The dashed line across the XY section indicates the position of the corresponding YZ reconstruction. Scale bar = 2 µm.

D. Confocal imaging (related to Figure 4H) of BMDMs expressing Akt-PH-mScarlet (magenta) and Mem-mNG(green), treated with PI3K pathway inhibitors and, stimulated with 200 ng/mL CSF1 for 2.5 min. Representative individual XY optical sections show the localization of Akt-PH-mScarlet (magenta) and Mem-mNG(green).The dashed line across the XY section indicates the position of the corresponding YZ reconstruction. Scale bar = 2 µm.

E. Dot plot showing the average life-time of the Akt-PH localization. n = 12 macropinosome. (Movie 3)

F. Confocal imaging of WT BMDMs stained with anti-PIP3 antibody (magenta) and phallodin (green). The cells were starved overnight, stimulated with 200 ng/mL CSF1 for 2.5 min, fixed and permeabilized. Scale bar = 5 µm.

**G.** Confocal imaging of BMDMs expressing 2xFYVE-mScarlet (magenta) and Mem-mNG(green), treated with or without PI3K pathway inhibitors and, stimulated with 200 ng/mL CSF1 for 2.5 min. Representative individual XY optical sections show the localization of Akt-PH-mScarlet (magenta) and Mem-mNG(green).The dashed line across the XY section indicates the position of the corresponding XZ or YZ reconstruction. Scale bar = 5 µm.

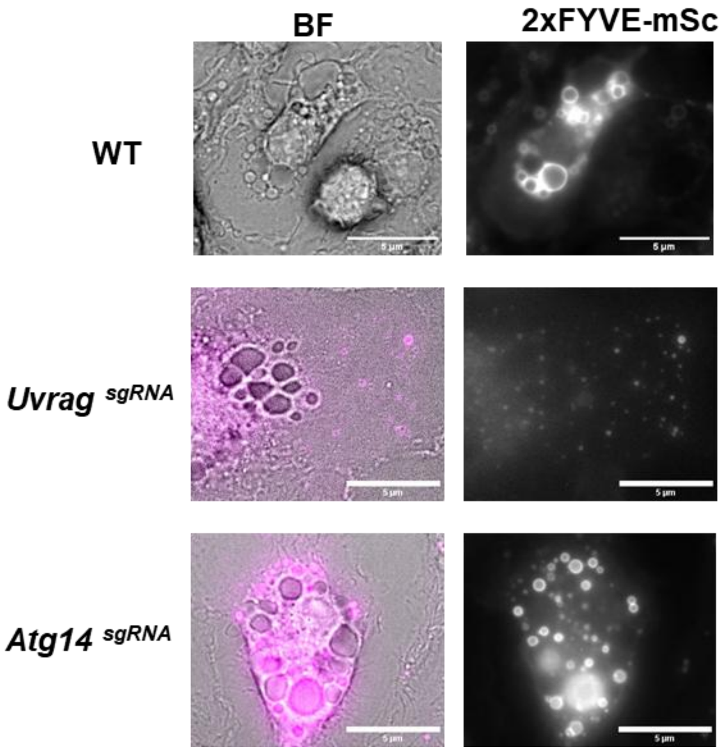


**Supplemental figure S4.**

Epifluorescence microscopic imaging of BMDMs expressing 2xFYVE-mScarlet (magenta). BMDMs were starved overnight, stimulated with 200 ng/ml CSF1 and imaged.Scale bar = 5 µm.

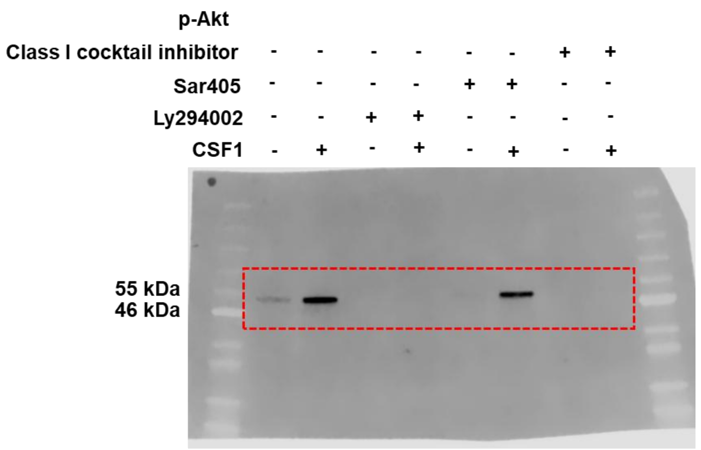


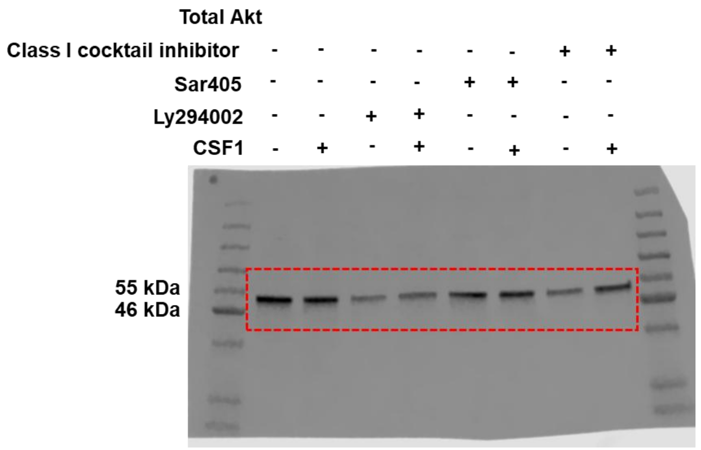


**Supplemental Figure S5.**

Unedited and uncropped western blots displayed in Figure 4B

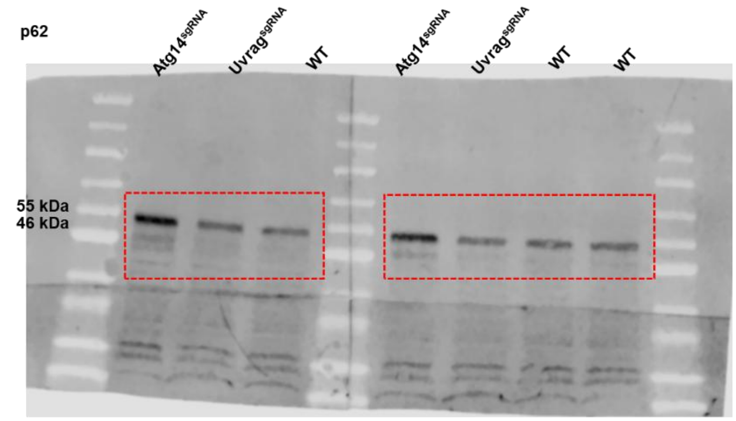


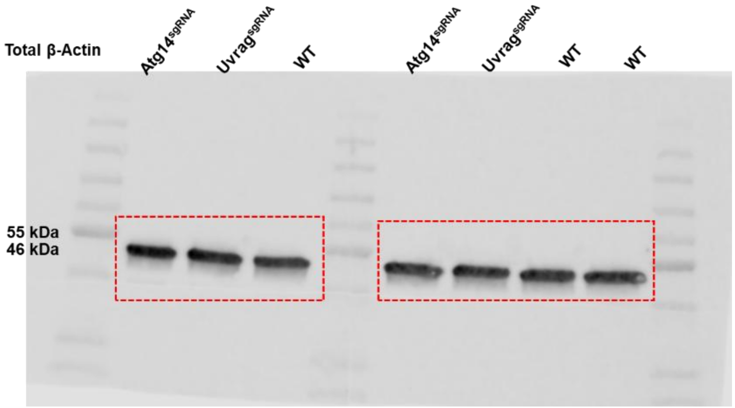


**Supplementary Figure S6**

Unedited and uncropped western blots displayed in Figure 6 B

### Movie legend

**Movie 1:** Brightfield microscopic movie of WT BMDMs related to Figure . Cells were starved overnight in DMEM containing 10% FBS and stimulated with CSF1. Cells were imaged in HBSS at 37°C using an Olympus IX83 microscope platform (Olympus, Shinjuku City, Tokyo, Japan) with a 60× oil-immersion objective (NA 1.42). Brightfield images were acquired every 15 s using a cooled CCD camera for a total duration of 8 min. The movie is displayed at 32 frames per second (fps). Scale bar = 2 µm.

**Movie 2:** Brightfield microscopic movie of $Uvrag^{sgRNA}$. BMDMs related to Figure . Cells were starved overnight in DMEM containing 10% FBS and stimulated with CSF1. Cells were imaged in HBSS at 37°C using an Olympus IX83 microscope platform (Olympus, Shinjuku City, Tokyo, Japan) with a 60× oil-immersion objective (NA 1.42). Brightfield images were acquired every 15 s using a cooled CCD camera for a total duration of 8 min. The movie is displayed at 32 frames per second (fps). Scale bar = 2 µm.

**Movie 3:** Brightfield microscopic movie of $Atg14^{sgRNA}$. BMDMs related to Figure . Cells were starved overnight in DMEM containing 10% FBS and stimulated with CSF1. Cells were imaged in HBSS at 37°C using an Olympus IX83 microscope platform (Olympus, Shinjuku City, Tokyo, Japan) with a 60× oil-immersion objective (NA 1.42). Brightfield images were acquired every 15 s using a cooled CCD camera for a total duration of 8 min. The movie is displayed at 32 frames per second (fps). Scale bar = 2 µm.

**Movie 4:** Brightfield microscopic movie of WT BMDMs related to Figure 4 C. Cells were starved overnight in DMEM containing 10% FBS and stimulated with CSF1. Cells were imaged in HBSS at 37°C using an Olympus IX83 microscope platform (Olympus, Shinjuku City, Tokyo, Japan) with a 60× oil-immersion objective (NA 1.42). Brightfield images were acquired every 15 s using a cooled CCD camera for a total duration of 9 minutes 45 second. The movie is displayed at 40 frames per second (fps). Scale bar = 1 µm.

**Movie 5:** Recruitment of PIP3 and PI(3)P during macropinosome formation. Live-cell fluorescence imaging of WT BMDMs expressing AktPH-mScarlet (green) and 2xFYVE-mScarlet (magenta)upon CSF-1 stimulation (related to Figure 4C). Cells were starved overnight in DMEM containing 10% FBS and stimulated with CSF1. Cells were imaged in HBSS at 37°C using an Olympus IX83 microscope platform (Olympus, Shinjuku City, Tokyo, Japan) with a 60× oil-immersion objective (NA 1.42). Images were acquired every 15 s using a cooled CCD camera for a total duration of 9 minutes 45 second. The movie is displayed at 40 frames per second (fps). Scale bar = 1 µm.

**Movie 6:** Brightfield microscopic movie of WT BMDMs related to Figure 4 E. Cells were starved overnight in DMEM containing 10% FBS and stimulated with CSF1. Cells were imaged in HBSS at 37°C using an Olympus IX83 microscope platform (Olympus, Shinjuku City, Tokyo, Japan) with a 60× oil-immersion objective (NA 1.42). Brightfield images were acquired every 15 s using a cooled CCD camera for a total duration of 9 minutes 45 second. The movie is displayed at 40 frames per second (fps). Scale bar = 1 µm.

**Movie 7:** Live-cell fluorescence movie showing an unclosed macropinocytic cup in WT BMDMs expressing AktPH-mScarlet (green) and 2xFYVE-mScarlet (magenta) upon CSF-1 stimulation (related to Figure 4E). Cells were starved overnight in DMEM containing 10% FBS and stimulated with CSF1. Cells were imaged in HBSS at 37°C using an Olympus IX83 microscope platform (Olympus, Shinjuku City, Tokyo, Japan) with a 60× oil-immersion objective (NA 1.42). Images were acquired every 15 s using a cooled CCD camera for a total duration of 9 minutes 45 second. The movie is displayed at 40 frames per second (fps). Scale bar = 1 µm.

**Movie 8:** Live-cell fluorescence movie showing the recruitment of AktPH in the cup following CSF1 stimulation. WT BMDM cells expressing LCK MemNG (green) and AktPH-mSC (magenta) were starved overnight in DMEM containing 10% FBS and stimulated with CSF1 for 2.5 mins. Cells were imaged in HBSS at 37°C using an Olympus IX83 microscope platform (Olympus, Shinjuku City, Tokyo, Japan) with a 60× oil-immersion objective (NA 1.42). Brightfield images were acquired every 15 s using a cooled CCD camera for a total duration of 5 minutes.. Scale bar = 2 µm.

**Movie 9:** Live-cell fluorescence movie showing the recruitment of AktPH in the cup following CSF1 stimulation. WT BMDM cells expressing LCK MemNG (green) and AktPH-mSC (magenta) were starved overnight in DMEM containing 10% FBS, treated with SAR405 and stimulated with CSF1 for 2.5 mins in the presence of SAR405. Cells were imaged in HBSS at 37°C using an Olympus IX83 microscope platform (Olympus, Shinjuku City, Tokyo, Japan) with a 60× oil-immersion objective (NA 1.42). Images were acquired every 15 s using a cooled CCD camera for a total duration of 5 minutes.. Scale bar = 2 μm.

**Movie 10:** Live-cell fluorescence movie showing the recruitment of AktPH in the cup following CSF1 stimulation. WT BMDM cells expressing LCK MemNG (green) and AktPH-mSC (magenta) were starved overnight in DMEM containing 10% FBS, treated with Class I inhibitor cocktail and stimulated with CSF1 for 2.5 mins in the presence of Class I inhibitor cocktail . Cells were imaged in HBSS at 37°C using an Olympus IX83 microscope platform (Olympus, Shinjuku City, Tokyo, Japan) with a 60× oil-immersion objective (NA 1.42). Brightfield images were acquired every 15 s using a cooled CCD camera for a total duration of 5 minutes.. Scale bar = 2 μm.

**Movie 11:** Lattice light sheet microscopy of WT BMDM related to figure 5.
